# Genetic mapping identifies Homer1 as a developmental modifier of attention

**DOI:** 10.1101/2023.03.17.533136

**Authors:** Zachary Gershon, Alessandra Bonito-Oliva, Matt Kanke, Andrea Terceros, Genelle Rankin, John Fak, Yujin Harada, Andrew F. Iannone, Millennium Gebremedhin, Brian Fabella, Natalia V. De Marco García, Praveen Sethupathy, Priya Rajasethupathy

## Abstract

Attention is required for most higher-order cognitive functions. Prior studies have revealed functional roles for the prefrontal cortex and its extended circuits to enabling attention, but the underlying molecular processes and their impacts on cellular and circuit function remain poorly understood. To develop insights, we here took an unbiased forward genetics approach to identify single genes of large effect on attention. We studied 200 genetically diverse mice on measures of pre-attentive processing and through genetic mapping identified a small locus on chromosome 13 (95%CI: 92.22-94.09 Mb) driving substantial variation (19%) in this trait. Further characterization of the locus revealed a causative gene, Homer1, encoding a synaptic protein, where down-regulation of its short isoforms in prefrontal cortex (PFC) during early postnatal development led to improvements in multiple measures of attention in the adult. Subsequent mechanistic studies revealed that prefrontal *Homer1* down-regulation is associated with GABAergic receptor up-regulation in those same cells. This enhanced inhibitory influence, together with dynamic neuromodulatory coupling, led to strikingly low PFC activity at baseline periods of the task but targeted elevations at cue onset, predicting short-latency correct choices. Notably high-*Homer1*, low-attentional performers, exhibited uniformly elevated PFC activity throughout the task. We thus identify a single gene of large effect on attention - Homer1 – and find that it improves prefrontal inhibitory tone and signal-to-noise (SNR) to enhance attentional performance. A therapeutic strategy focused on reducing prefrontal activity and increasing SNR, rather than uniformly elevating PFC activity, may complement the use of stimulants to improve attention.

## INTRODUCTION

Attention is the process of directing cognitive resources to particular stimuli and is a pre-requisite for higher-order cognition, such as short-term memory. Nonetheless, there is still much debate in the field on how to conceptualize attention. Many have conceived of it as a collection of distinct, but related processes; that is, constructs such as orienting, expectancy, and stimulus differentiation are engaged for selective attention, yet all of those processes are then necessary to report unpredictable events over a prolonged period of time, i.e. sustained attention^1–3^. This complexity is deepened as attention can be constant or fluctuating, occurring on slow or rapid time-scales, and can be broadly distributed in the brain but also highly specific to particular stimuli^4–8^. Despite these distinctions, pre-attentive processing to filter out irrelevant stimuli is an early component thought to be needed for many if not all downstream aspects of attention. Years of foundational research have highlighted the importance of enhanced prefrontal activity in mediating pre-attentive processing and attentional control. It is also well appreciated that the PFC interfaces with a broader neuroanatomically distributed network to enable attention. For instance, long-range recurrence via thalamus^9,10^, and neuromodulation via adrenergic, dopaminergic and other systems^11–15^, are thought to be key mediators of attention. Indeed, many ADHD medications target these circuits^16^. However, there are important limitations in our understanding of the specific circuits, cell-types, and underlying molecular pathways involved in this cognitive process. More importantly, we lack an understanding of which nodes in these complex pathways are most critical, which if identified, can inform more unifying models and therapeutic strategies for attentional processing.

In the past, unbiased genetic mapping approaches enabled the identification of single genes with large contributions to a behavioral trait^17–20^. Further investigations of these genes identified key cell types and circuits that led to unifying cellular models of behavior. Toward this goal, we recently performed genetic mapping in mice and identified a single gene with large contributions to short-term memory^21^. Building on the fruition of this previous work, in this study we focused on identifying genetic contributions to attention.

In our prior experiences, genetic mapping in outbred mice can be most successful if the screening behaviors are simple, innate, and robustly quantifiable. Since traditional tasks of attention require extensive training (often 3-6 weeks), reward-associations, and other potential confounds for genetic mapping, we selected and optimized an assay for an innate pre-attentive processing behavior, i.e., prepulse inhibition of startle response (PPI). This behavioral process of PPI refers to the ability of an animal to suppress a startle response to a sudden strong stimulus when preceded by a weaker stimulus, which is thought to reflect the process of neural filtering of redundant or irrelevant stimuli while enhancing subsequent goal-directed processing of salient aspects of the environment^22^. Extensive prior work has characterized PPI as a pre-attentive process^23–27^ and additional studies have linked it to downstream measures of attention in rodents and humans^28–34^. We thus screened a cohort of genetically diverse mice on this pre-attentive processing task and identified a genetic locus of large effect on chromosome 13 linked to variation in this trait. Further characterization of the locus on more targeted measures of attention revealed a causative gene, Homer1, encoding a synaptic protein, where down-regulation of its short isoforms in PFC during development enhances multiple measures of attentional performance in adult mice. Notably, low-*Homer1a*, high-attention mice were associated with increased prefrontal receptivity to inhibition, dynamic task-associated scaling of neural activity, and increased signal to noise (SNR) during task performance. We thus identify a single gene with large effect on attention – Homer1 – and link this with increased SNR, rather than overall magnitude, of PFC activity as an important component of attention.

## RESULTS

### Identification of a QTL linked to pre-attentive processing

The Diversity Outbred (DO) resource is a mouse population that is derived from eight founder strains, whose genetic diversity, including SNP density and allelic heterozygosity, is comparable to that of the human population, providing a platform for high-resolution genetic mapping (Figure 1A). Based on our previous work^21^ characterizing the DO founders, and cohorts from the 19^th^ and 25^th^ generations, we determined that successful mapping of genetic loci to behavioral traits would require 1) An automated and robust behavioral assay requiring minimal training, thus narrowing the observed variance to genetic and task-associated features, and 2) Approximately 200 mice to detect a quantitative trait locus (QTL) that shifts the trait mean by 1 standard deviation at 95% confidence. Since traditional tests of attention require weeks of training, we selected and optimized a simple and robust assay to test an innate pre-attentive processing behavior, referred to as pre-pulse inhibition (PPI). PPI is the ability to suppress an innate startle response, which reflects a process of filtering irrelevant cues to enhance goal-directed selection. Notably, while the startle response is considered a “bottom-up” process^28^, the inhibition of startle is a “top-down” process^32^ that has been linked to downstream measures of cognition, including selective^34^ and sustained^33^ attention. Although PPI can also be confounded by changes in anxiety and motor response (which we tested post hoc), we chose this task as a fast, high-throughput, sensitive screen for pre-attentive processing, which we could then follow up with more targeted tests for attention.

**Figure 1.**
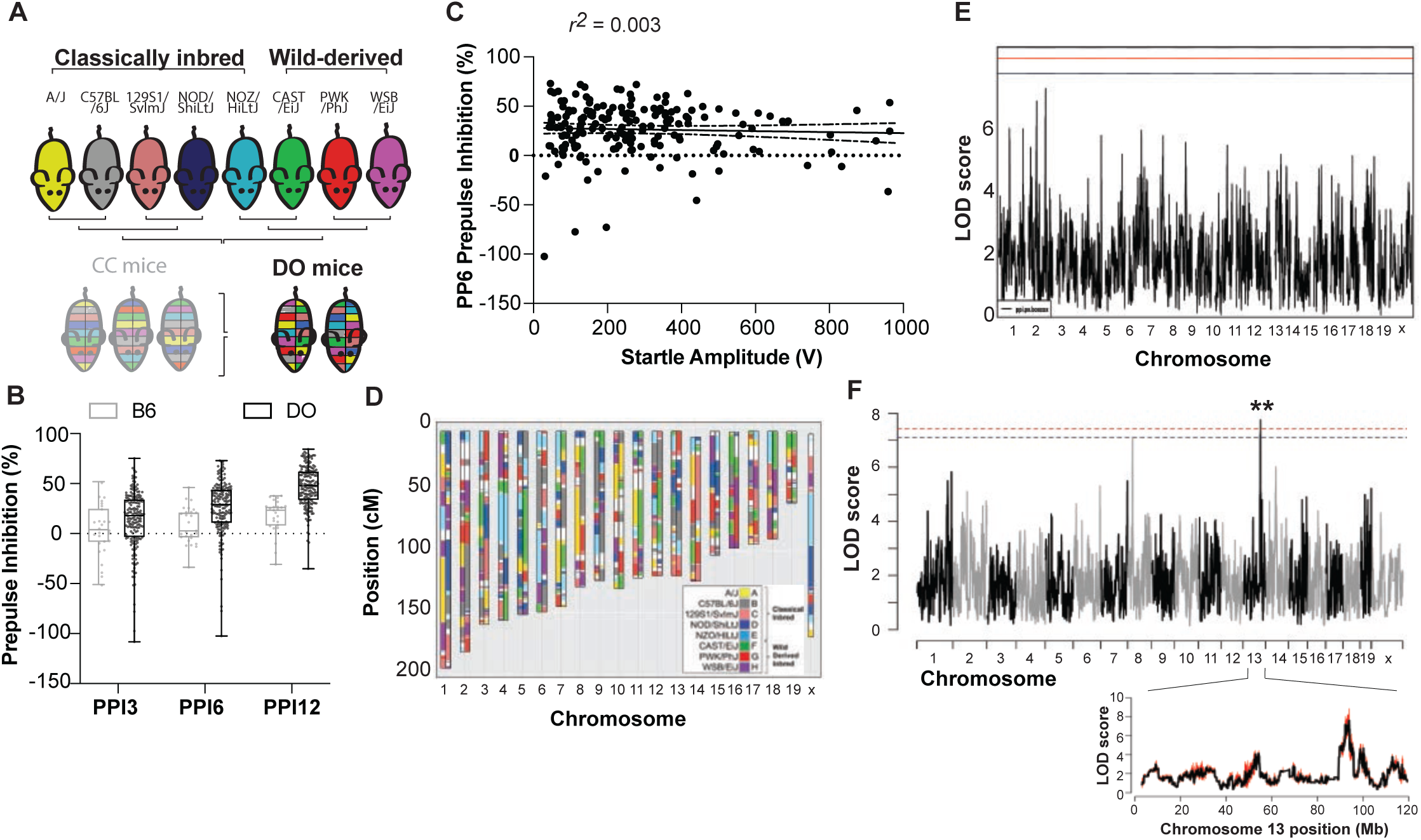
Identification of a QTL associated with pre-attentive processing. (**A**) Outbreeding scheme to generate the DO mice. (**B**) Pre-attentive processing performance (assayed by PPI) in B6 (n=27) and DO (n=176) mice measured as percent of startle response inhibited at 3 different pre-pulse intensities: 3, 6, and 12 dB above background (PPI3, 6, and 12, respectively). Boxes indicate 2nd and 3rd quartiles with median and range. (**C**) Correlation in DO mice (n=176) between startle response, measured as the magnitude of startle amplitude (V), and PPI, measured as percent inhibition, at 6 dB above background (PP6, r^2^=0.003). (**D**) Haplotype reconstruction of a representative DO mouse from the 25th generation of the population. Colors correspond to the founder lines (shown in legend) for which the genomic contribution is attributed at each depicted locus. (**E**) QTL mapping analysis of startle response (by R/qtl2) shown as a Manhattan plot of startle response. Blue and red lines indicate confidence thresholds, blue: 90%, red: 95%. (**F**) Top: QTL analysis (by miQTL) for PPI at 6 dB above background (PPI6). Confidence thresholds after 50 imputations of genotype, blue: 90%, red: 95%. Genome-wide p<0.01. Bottom: Mapping analyses performed using R/qtl2 (black) and miQTL (red) revealing minimal fluctuation in LOD score across imputations (overlapping bands).

We tested 191 mice for performance in PPI. Briefly, for each DO mouse we measured the startle response to a 120 dB tone as well as the percent inhibition of this startle when preceded by a weaker 3, 6, or 12 dB tone (PPI3, PPI6, PPI12). We first confirmed that the phenotypic variability of the DO greatly surpassed that of the C57BL/6J (B6) classical inbred line, as would be expected from the underlying genetic variation (Figure 1B). We excluded 15 mice that exhibited greater PPI3 than PPI12, suggesting potential hearing impairment. With the remaining mice, we found no significant correlations between PPI and startle response or body weight (Figures 1C and S1A-D).

We next genotyped the 176 DO mice using the GigaMUGA platform (114,184 loci had variability in our cohort). Founder haplotype reconstructions were performed with a hidden Markov model^21,35^, which showed extensive allelic heterozygosity genome-wide (Figure 1D) and we observed approximately equal founder contributions across our cohort suggesting minimal allelic loss. We performed QTL mapping for PPI using R/qtl2^23^ and identified a single large effect genetic locus (19% of behavioral variance explained) on chromosome 13 with genome-wide significance of p ≤0.01 (Figure S1E; LOD score for PPI6 = 8.22, 95%CI: 92.22-94.09 Mb). These mapping effects were not due to individual differences in the underlying innate startle response (Figure 1C), nor was there any QTL detected when mapping to startle scores (Figure 1E). The chromosome 13 QTL for PPI6 was also confirmed to be statistically significant using a second mapping approach, miQTL (Figure 1F). QTL mapping of PPI3 and PPI12 did not reveal any loci that surpassed significance thresholds, but a suggested peak for PPI3 indeed mapped to the same Chr 13 QTL (Figure S1E; LOD score = 5.88, 95%CI: 90.51-94.09 Mb), supporting the functional significance of this locus.

Next, to further increase confidence in this locus we performed an allele effect analysis (Methods) and found that the B6 haplotype (henceforth referred to as Chr13QTL^B6^) was associated with high performance while the WSB/EiJ haplotype (henceforth referred to as Chr13QTL^WSB^) was associated with low performance (Figure 2A-B). We then asked whether specific recombinant inbred Collaborative Cross (CC) lines, which have the same multi-parent origins as the DO (Figure 2C) and possess either Chr13QTL^B6^ or Chr13QTL^WSB^ would separate into high and low performers, respectively. We analyzed the genomes of existing CC lines and selected two that were homozygous for our desired Chr13QTL^B6^ (CC083) or Chr13QTL^WSB^ (CC025) haplotypes (Figure 2D), while maintaining distinctive mosaic representations of the founder genomes at other loci. We compared PPI performance between CC083 and CC025 and found that CC083 have significantly greater PPI than CC025 (Figure 2E). As with the DO, this finding was not explained by differences in peak startle or body weight (Figures S2A-D), nor was it due to differences in gross motor activity, motor coordination or hearing sensitivity (Figures 2F-H). These data indicate that genetic variation at the Chr13 locus, specifically WSB vs B6 genotype, drives significant variation in pre-attentive processing.

**Figure 2.**
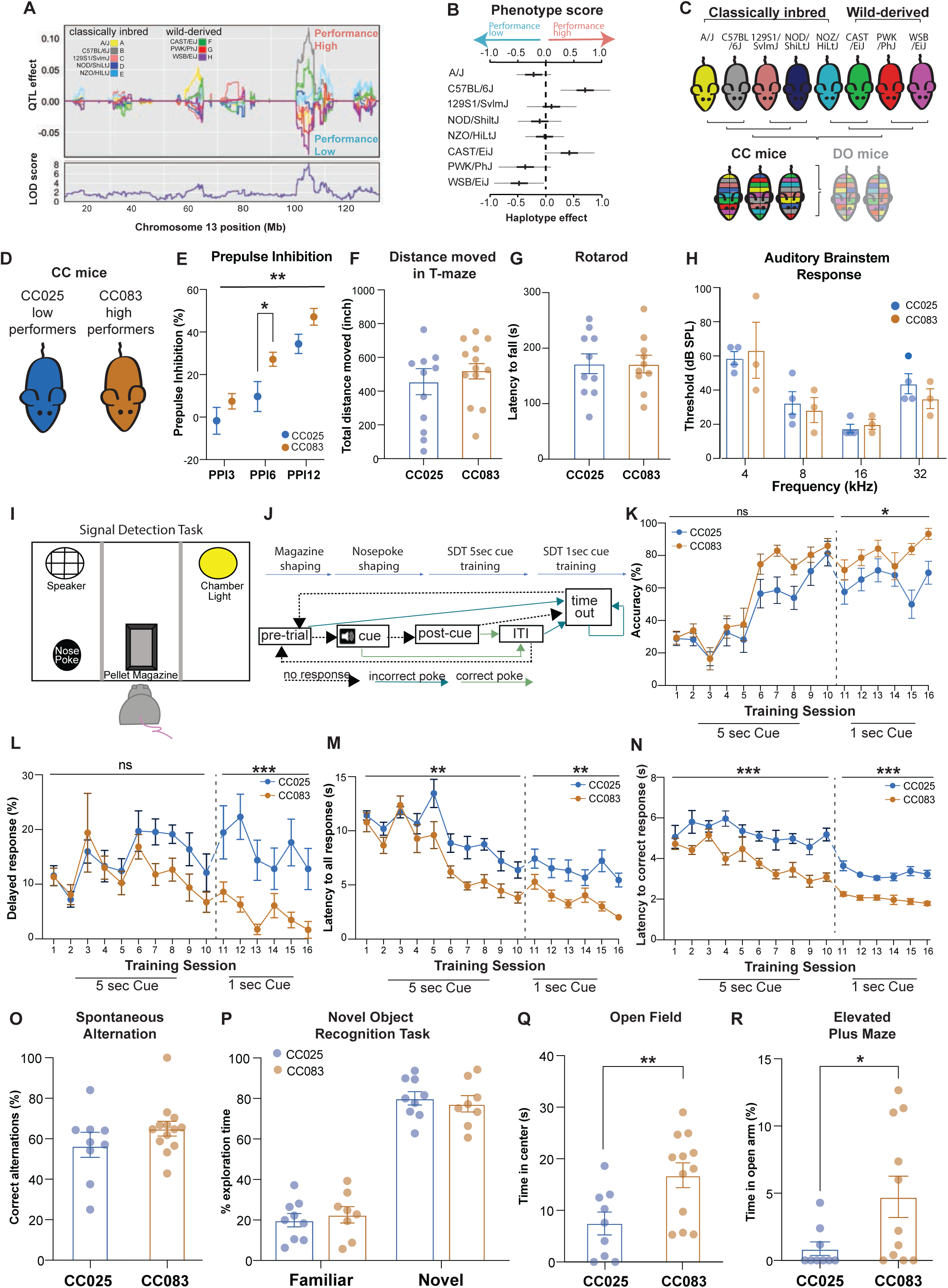
Chr13 QTL mediates variation in attentional performance. (**A**) Effect of each founder allele on PPI performance along Chromosome 13, as measured by founder coefficients from the linkage model. Coefficients diverge substantially at peak QTL. Logarithm of odds (LOD) score at each chromosomal position shown. (**B**) Haplotype representation at the Chromosome 13 locus and corresponding z-scored phenotypes of each founder strain, quantified as mean ± 95% confidence intervals. (**C**) Outbreeding scheme to generate the CC mice. (**D**) Cartoon of the CC025 (Low performers, blue) and CC083 (High performers, tan) used in subsequent experiments. (**E**) PPI3, 6, and 12 values for CC083 (n=27) and CC025 (n=24) mice. Two-way ANOVA p=0.003 for CC line main effect followed by Holm-Sidak’s test for multiple comparison p_PPI6_=0.05. (**F**) Gross motor activity measured in CC025 (n=11) and CC083 (n=11) mice as total distance moved (inch) in a T-maze apparatus during a 6-min test. (**G**) Motor coordination measured in CC025 (n=10) and CC083 (n=10) as latency (s) to fall from the rod in the Rotarod test averaged across 4 consecutive trials. (**H**) Auditory brainstem response measured as minimum thresholds in CC025 (n=4) and CC083 (n=3) as sound pressure level (dB) in response to increasing frequencies (4, 8, 16, 32 kHz). (**I**) Schematic of operant wall of arena used for signal detection task (SDT). (**J**) Schematic of SDT protocol. (**K-N**) Performance of CC025 (n=10 for 5 sec cue and n=9 for 1 sec cue) and CC083 (n=10 for 5 sec cue and 1 sec cue) mice during SDT across sessions, showing (**K**) accuracy (correct response) percentage (repeated-measures two-way ANOVA p= 0.02 for CC line main effect in 1 sec cue sessions), (**L**) delayed response percentage (repeated-measures two-way ANOVA p<0.001 for CC line main effect in 1 sec cue sessions), (**M**) mean latency from cue to first response within all trials (repeated-measures two-way ANOVA for p=0.009 and p=0.002 for CC line main effect in 5 sec cue and 1 sec cue sessions, respectively), and (**N**) mean latency from cue to first response within correct trials (repeated-measures two-way ANOVA p<0.001 for CC line main effect in both 5 and 1 sec cue sessions). (**O**) Working memory performance assessed in a T-maze apparatus for CC025 (n=9) and CC083 (n=13) mice, measured as the percent of correct alternations (Methods). (**P**) Short-term memory tested by a novel object recognition test in CC025 (n=9) and CC083 (n=10) mice, measured as time spent exploring the novel object *vs* the familiar one and expressed as percentage of total exploration time during a 5-min test. Two-way ANOVA showed significant main effect for novelty (p<0.001), but not for CC line. (**Q,R**) Anxiety-like behavior measured as (**Q**) time, in seconds, spent in the center of an open field arena during a 5-min test in CC025 (n=9) and CC083 (n=12). Welch-corrected t-test showed a significant difference between CC lines (p=0.01) and (**R**) percentage of time spent in the open arm of an elevated plus maze during a 5-min test in CC025 (n=11) and CC083 (n=10) mice. Welch-corrected t-test showed a significant difference between lines (p=0.03). Data in **D**-**R** are expressed as mean ± SEM.

To more directly test the role of this Chr 13 QTL in attention, we studied CC083 and CC025 mice in a more targeted assay for attention, an operant signal detection task. Here, mice are trained to nose-poke in response to a 5 second auditory cue within 10 second of cue onset to receive a food reward. Once the mice have sufficiently learned the task (Methods), their attentional load is then challenged by decreasing the length of the cue to 1 sec and reducing the response window (Figures 2I-J). Similar signal detection tasks have been widely used to assay attention^36–38^. They provide multiple metrics to track attention including accuracy, response latency, and trial omissions. During the initial 5 sec cue training, there were no significant differences in learning the task, but CC083 mice were already exhibiting fast latency responses, and after increased attentional load during the 1 sec trials, the CC083s significantly outperformed the CC025s in all of measures of attention including accuracy (percentage of correct responses), proportion of delayed responses, latency to all responses, and, most significantly, latency to correct responses (Figures 2K-N). Notably, the strains did not differ in other cognitive, or social measures that we tested (Figures 2O-P and S2E). We did observe differences in measures of anxiety-related behavior (Figure 2Q-R), which requires further consideration given important dependencies between anxiety and attention (although of note, in later experiments, when manipulating only the causal gene at this locus, no significant differences in anxiety-like behavior was observed, Figures 4U-V). Together, these data indicate that genetic variation at the chromosome 13 locus drives differences in attentional performance.

### Chr 13 QTL effects on attention are driven by *Homer1*

We next sought to understand which gene(s) within the locus was driving the changes in attentional performance. We performed bulk RNA sequencing in DO high and low performers, focusing on the prefrontal cortex (PFC) because of its central role in attentional processing, but also including related brain areas such as mediodorsal thalamus (MD), and the ventral tegmental area (VTA). We found that samples stratified by performance in PFC and MD, but not in VTA (Figure S3A), leading us to ask which genes within the Chromosome 13 locus (Figure S3B) were differentially expressed in MD or PFC between high and low performers. Of all locus genes, only *Homer1* was significantly differentially expressed, with substantial downregulation in PFC in high performers (Figure 3A, adjusted p<0.001). *Homer1* has several transcript variants due to alternative splicing (Figure S3C)^39^, and thus we assessed whether differential expression was uniform across splice isoforms. Strikingly, only the short, activity-dependent isoforms, *Homer1a*^40,41^ and *Ania3*^39^, were differentially expressed between DO high and low performers (Figure 3B, p(*Homer1a*)=0.003, p(*Ania3*)=0.007, two-way ANOVA with post hoc Holm-Sidak test for multiple comparisons). Furthermore, bulk RNAseq from high (CC083) and low (CC025) performing CC lines also confirmed significant differences in *Homer1* and associated gene ontology terms relating to excitatory neurotransmission (Figure 3C). As with the DO mice, the differential *Homer1* expression in CC mice (Figure 3D) was driven by downregulation of *Homer1a* and *Ania3* in the high performing CC083s (Figure 3E, 2-way ANOVA p<0.001, Holm-Sidak test for multiple comparisons p(*Homer1a*)<0.001, p(*Ania3*)<0.001).

**Figure 3.**
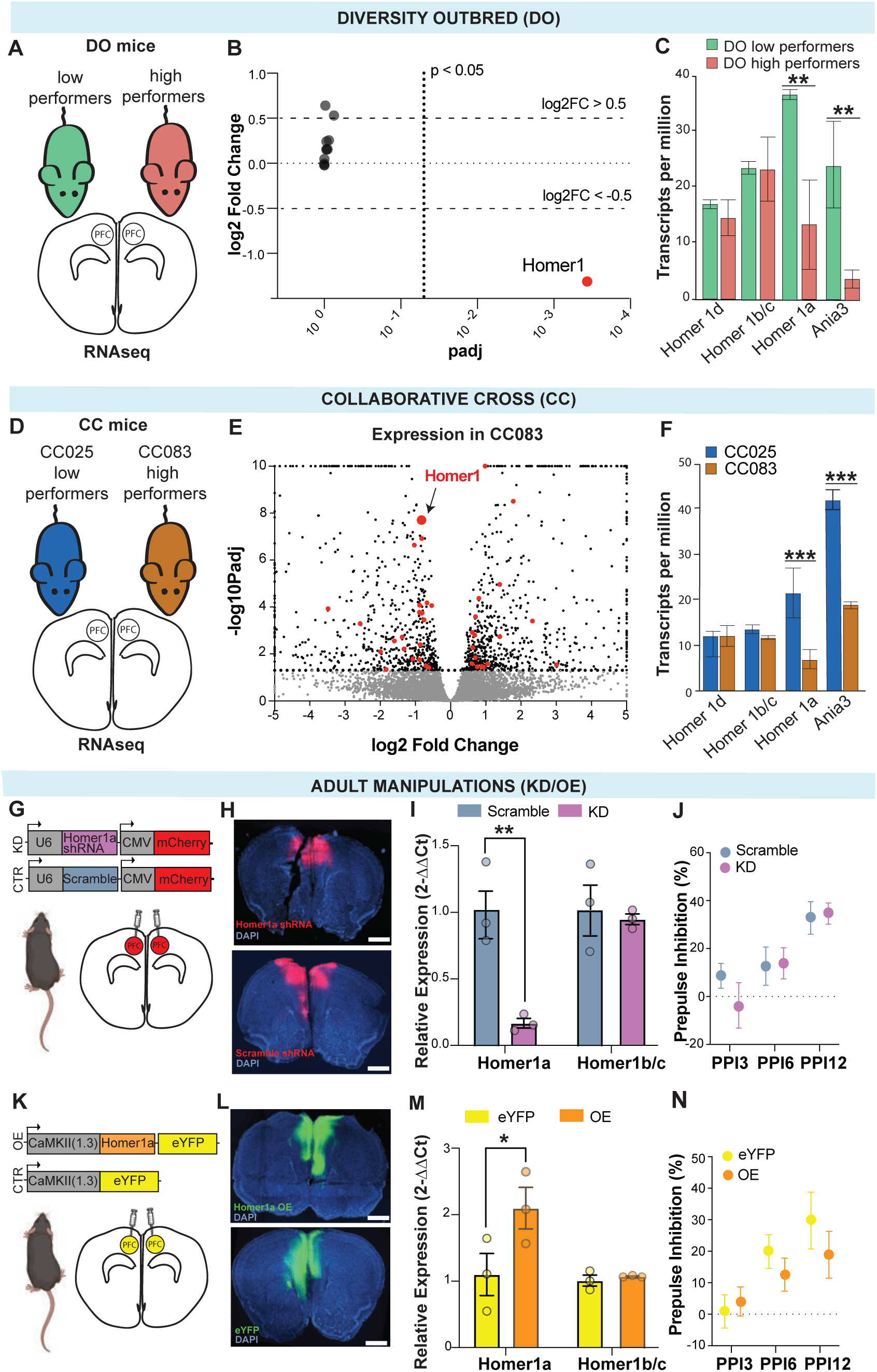
Chr13 QTL effects map to Homer1, but adult manipulations have no behavioral phenotype. (**A**) Schematic of prefrontal cortex (PFC) dissection region for RNAseq in DO high (pink) and low (green) performers. (**B**) Volcano plots of differential expression between DO high (pink) and low (green) performers for all locus genes (n=3 per group) from bulk PFC RNAseq. Dashed lines indicate significance thresholds (adjusted p=0.05 and log2FC=0.5 or =-0.5). Only *Homer1* crosses both thresholds (red). (**C**) Expression levels of *Homer1* isoforms in PFC from DO high and low performers (n=3 per group), significant differential expression of *Homer1a* and *Ania3*, p<0.01 by two-way ANOVA with post hoc Holm-Sidak’s test. (**D**) Schematic of PFC dissection region for RNAseq in CC high (CC083, tan) and low (CC025, blue) performers. (**E**) Volcano plot showing differential gene expression for CC083 (high performers) relative to CC025 (low performers) mice after DESeq2. X- and Y-axis denote log2 fold change and −log p value distribution, respectively. Red dots indicate genes with ontologies relating to *Homer1*’s function in excitatory neurotransmission and postsynaptic structure & activity. (**F**) Expression levels of *Homer1* isoforms in PFC from CC high and low performers (n=3 per group), p<0.001 by two-way ANOVA with post hoc Holm-Sidak’s test for multiple comparisons. (**G**) Schematic of constructs and injection location (PFC) for knockdown (KD, purple) and control (Scramble, blue) in adult B6 mice. (**H**) Validation histology performed 8 weeks after bilateral injection of AAV-U6-Homer1a shRNA-CMV-mCherry knockdown virus (upper panel) and AAV-U6-Scramble-CMV-mCherry control virus (lower panel) into PFC showing viral transduction in the target area (DAPI, blue; mCherry, red), scale bars: 1000µm.(**I**) *Homer1a* and *Homer1b/c* expression levels (relative to controls) in PFC samples dissected from KD (n=3) and control (n=3) mice measured by qPCR, (two-way ANOVA showed significant main effects for shRNA construct, p=0.007, and *Homer1* isoform expression, p=0.02, as well as a significant interaction between those variables, p=0.02; post hoc Holm-Sidak’s test for multiple comparisons shows a significant difference in *Homer1a* expression, p=0.003). (**J**) PPI in KD (n=15) and Scramble (n=15) mice measured as percent inhibition at 3 prepulse intensities: 3, 6, and 12 dB above background (PPI3, 6, and 12, respectively). (**K**) Schematic of constructs and injection location (PFC) for overexpression (OE, orange) and control (eYFP, yellow) in adult B6 mice. (**L**) Validation histology performed 8 weeks after bilateral injection of AAV-CaMKII(1.3)-eYFP overexpression virus (upper panel) and AAV-CaMKII(1.3)-eYFP control virus (lower panel) into PFC, showing viral transduction in the target area (DAPI, blue; eYFP, green), scale bars: 1000µm.(**M**) *Homer1a* and *Homer1b/c* expression levels (relative to controls) in PFC samples dissected from OE (n=3) and control eYFP (n=3) mice measured by qPCR (two-way ANOVA showed significant main effects for expression construct, p=0.04, and *Homer1* isoform expression, p=0.04; post hoc Holm-Sidak’s test for multiple comparisons shows a significant difference in *Homer1a* expression, p=0.03). (**N**) PPI in OE (n=11) and control eYFP (n=10) mice measured as percent inhibition at 3 pre-pulse intensities: 3, 6, and 12 dB above background (PPI3, 6, and 12, respectively). Data in **C**, **F**, **I**, **J**, **M**, and **N** are expressed as mean ± SEM.

Since *Homer1a* is better characterized and conserved than *Ania3*^39,42^, we next assessed whether *Homer1a* manipulations could drive behavioral changes in attentional performance. To knock down *Homer1a*, we designed and tested AAV-based short-hairpin RNAs (shRNAs) to target the *Homer1a* isoform *in vitro* and selected the most effective shRNA (Figures S3D-E) for bilateral PFC injections *in vivo* and behavioral testing (Figures 3G-J & Figure S3F). To overexpress *Homer1a,* which has endogenous expression primarily in excitatory pyramidal neurons, we cloned the *Homer1a* coding sequence into an AAV-based CaMKII-eYFP vector (Figure S3G) for bilateral PFC injection and behavioral testing (Figures 3K-N and S3H). To our surprise, we did not observe any significant behavioral effect for either the knockdown or overexpression experiments (Figures 3J, 3N, S3F, & S3H). To account for potential functional redundancy of Homer1a through Ania3, we performed bilateral PFC injections of the AAV shRNA targeting *Homer1a* pooled together with an AAV-based shRNA for *Ania3* (Figure S3I-J), which we validated *in vitro* (Figures S4A-B), and also saw no significant behavioral effect (S3K-L).

To assess whether the effects of *Homer1a* may be developmental in origin, based on prior work on germline knockouts^43–46^, we profiled the expression of *Homer1a*, *Ania3*, and *Homer1b/c* in CC083 and CC025 mice across postnatal development (Figure 4A). We found that, the expression of *Homer1a* and *Ania3,* but not that of Homer1b/c, diverged between the CC lines as early as p14-p21 (Figure 4B; two-way ANOVA p=0.02), suggesting possible developmental roles in regulating attentional processing. To test this hypothesis, we knocked down *Homer1a* and *Ania3* during early developmental stages (p14-p21) to evaluate the effect on adult behavior by bilaterally injecting the pooled *Homer1a* and *Ania3* shRNA AAVs into the PFC of neonatal B6 pups (Figure 4C; referred to as KD_dev_). Despite the developmental *Homer1a* knockdown being less effective than the adult manipulation (~80% in adults and ~60% in pups; Figures 3I & S4C), we observed significant improvement in measures of pre-attentive processing (PPI, Figure 4D).

**Figure 4.**
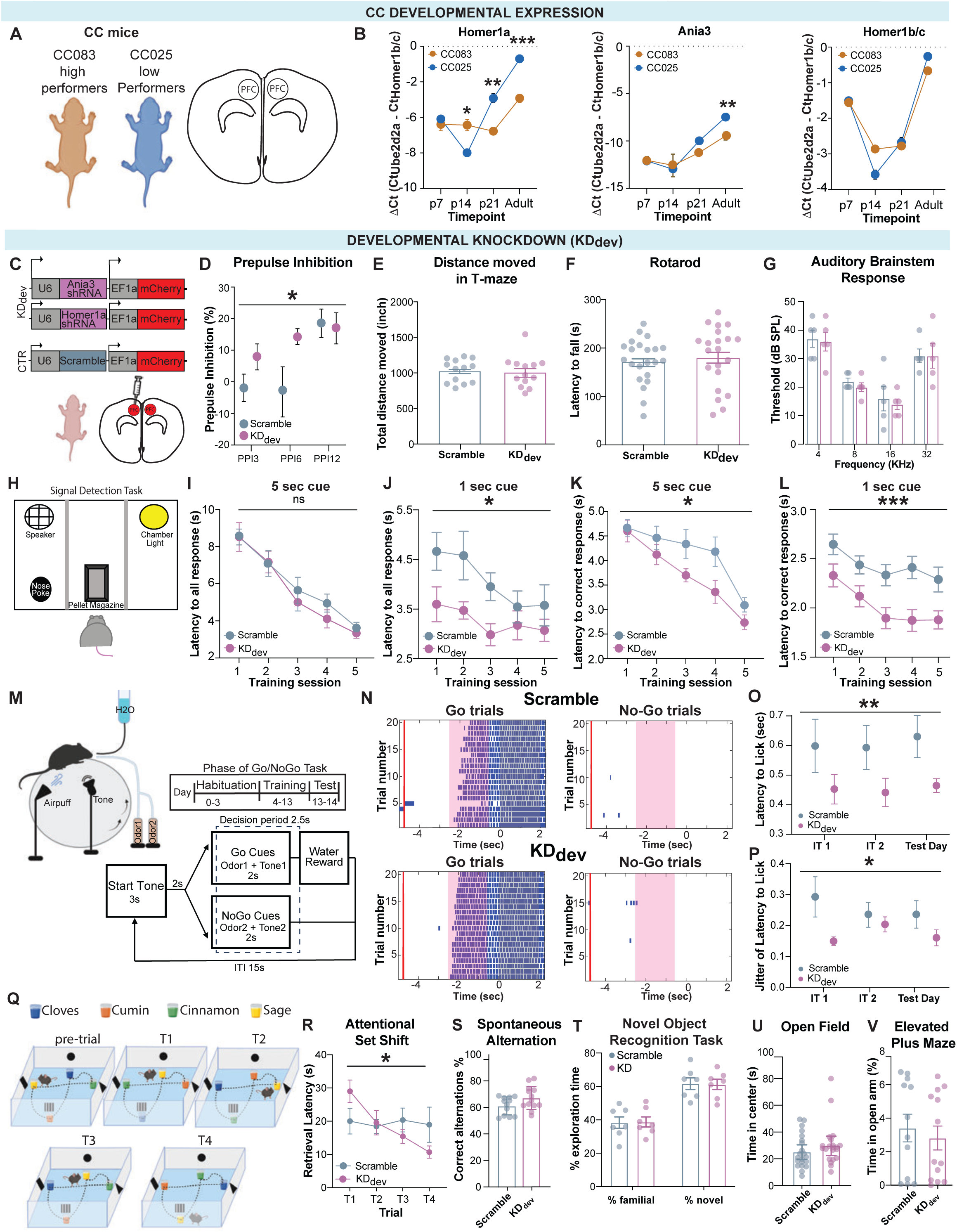
Homer1a and Ania3 are developmental modifiers of attention. (**A**) Schematic of PFC dissection region in CC high (CC083, tan) and low (CC025, blue) performers for qPCRs across postnatal development. (**B**) PFC expression of *Homer1a*, *Ania3*, and *Homer1b/c* in CC083 and CC025 mice at p7, p14, p21, and in adult by qPCR (n=3 per strain per timepoint), significant differences for *Homer1a* by two-way ANOVA with post hoc Holm-Sidak’s test, p=0.02 at p14, p=0.002 at p21, and p<0.001 at adult; and for *Ania3* p=0.002 at adult. (**C**) Schematic of constructs and injection location (PFC) for knockdown (KD_dev_, purple) and control (Scramble, blue) in neonatal B6 mice. (**D**) PPI in Scramble (n=13) and KD_dev_, (n=15). Significant differences between groups by two-way ANOVA (p=0.04). (**E**) Gross motor activity measured as distance moved (inch) by Scramble (n=15) and KD_dev_ (n=15) during a 6-min spontaneous alternation test in a T-maze apparatus. (**F**) Motor coordination in the Rotarod test for Scramble (n=22) and KD_dev_ (n=21), measured as latency (s) to fall from the rod averaged across 4 consecutive trials. (**G**) Auditory brainstem response measured as minimum thresholds in Scramble (n=5) and KD_dev_ (n=5), as sound pressure level (dB) in response to increasing frequencies (4, 8, 16, 32 kHz). (**H**) Schematic of operant wall of arena used for signal detection task (SDT). (**I-L**) Performance during SDT across training sessions, shown as (**I-J**) mean latency from cue to first response within all trials (p_1sec_=0.04) and (**K-L**) mean latency from cue to first response within correct trials (p_5sec_=0.03; p_1sec_<0.001) for 5 sec cue (Scramble n=13 and KD_dev_ n=13, **I** and **K**) and 1 sec cue (Scramble n=9 and KD_dev_ n=11, **J** and **L**). Significant differences between groups measured by repeated-measures two-way ANOVA. (**M**) Schematic of Go/No-Go task setup (left) and training protocol (right). Below is the task structure for interleaved training days testing day. (**N**) Raster plots of licking for the Go/No-Go task for representative Scramble (above) and KD_dev_ (below) mice. Go (right) and No-Go (left) trials were interleaved during testing but are depicted separately. Time 0 is plotted as the end of the decision period. The red bar shows the end of the start tone, pink shading notes the time when cues are delivered, and licks are plotted as blue ticks. (**O**) Quantification of the latency to first lick within the decision period of Go trials. Each point is the average latency to first lick for the first 10 Go trials per animal (p=0.005, n=8 per group, significant main effect between groups by two-way ANOVA). (**P**) Quantification of the latency to first lick jitter. Jitter was quantified as the standard deviation of first lick latencies across the first 10 Go trials (significant main effect between groups by two-way ANOVA showed a significant main effect between groups, p=0.01, n=8 per group). (**Q**) Schematic of the Attentional Set Shift set up and experiment protocol. (**R**) Latency (s) to retrieve the chocolate pellet measured in Scramble (n=14) and KD_dev_, (n=13) mice during the 4 trials of the Attentional Set Shift test. Significant interaction between trial and group, p=0.04, by repeated-measures two-way ANOVA. (**S**) Working memory performance assessed in a T-maze apparatus for Scramble (n=12) and KD_dev_, (n=13) mice, measured as correct alternations performed, expressed as a percentage total alternations. (**T**) Short-term memory tested by a novel object recognition test in Scramble (n=7) and KD_dev_, (n=7) mice, measured as time spent exploring the novel object *vs* the familiar one and expressed as a percentage of total exploration time during a 5-min test. Two-way ANOVA showed significant main effect for novelty (p<0.001), but not for line. (**U,V**) Anxiety-like behavior measured as (**U**) time (in seconds) spent in the center of an open field arena during a 5-min test in Scramble (n=21) and KD_dev_, (n=21) mice, and (**V**) time spent in the open arm (%) of an elevated plus maze during a 5-min test in Scramble (n=12) and KD_dev_, (n=13) mice. Data in **B**, **D**-**G**, **I**-**L**, **O**-**P**, **R**-**V** are expressed as mean ± SEM.

We next tested the effects of developmental *Homer1a/Ania3* knockdown on multiple, widely-used assays for attention in adult mice including: 1. An operant SDT (Figures 4H-L, S4H-I), 2. a Go/No-Go task (Figures 4M-P, S4J-K), 3. a head-fixed multi-modal SDT (Figure S4L-N) and 4. an attentional set shift task^47^ (Figure 4Q-R). In all cases, we observed that mice with developmental prefrontal *Homer1/Ania3* knockdown performed significantly better on measures of attention than the corresponding controls. For instance, on the operant SDT task, when comparing response latencies on correct trials, the most sensitive measure of attentional performance, KD_dev_ mice exhibited significantly faster response latencies than controls, particularly on correct trials, that persisted throughout the extent of both cue length phases (Figures 4I-L; repeated-measures two-way ANOVA p(5sec cue)=0.04, p(1sec cue)<0.001). Furthermore, in a head-fixed Go/No-Go task where mice were trained to respond to one paired tone/odor cue and inhibit response to a different paired tone/odor cue (Figure 4M), KD_dev_ mice responded faster (Figure 4N-O) and more reliably (Figure 4P) than scramble controls. Notably, the magnitude of these effect sizes were substantial, for instance, with mean differences in response latency between groups of ~400ms on the operant SDT task (~2.3s for Scramble controls vs 1.9s for KD_dev_, p<0.001 by two way ANOVA, Figure 4L) and ~150ms for the head-fixed Go/No-Go task (~650ms for Scramble controls vs 500ms for KD_dev_, p<0.01 by two way ANOVA, Figure 4O, and appreciable qualitatively in the raw lick rasters in Figure 4N). These effects between groups were not present prior to learning, and there were no obvious differences in overall ability to learn the tasks (Figures S4I & S4K).

In a head-fixed multi-modal signal detection task, where mice only had to respond to the presence of a paired tone/odor cue (Figure S4L-N), KD_dev_ mice responded only slightly faster but displayed strikingly less variability in their response latency than the scramble control mice (Figures S4M-N). Finally, using an odor-based attentional set shift task we found that KD_dev_ mice displayed significantly shorter latencies to retrieve the food reward than control mice (Figure 4Q-R), despite having a similar baseline exploratory activity (measured as time spent exploring the four odors during pre-trial when the reward was not present: Scramble 92.2 ± 8.8 and KD_dev_ 100.1 ± 12.7), or bias between odors (measured as % time spent exploring each odor / total exploration time (cloves: Scramble 27.5 ± 3.4, KD_dev_ 23.3 ± 2.7; cumin: Scramble 20.9 ± 1.9, KD_dev_ 21.9 ± 3.2; sage: Scramble 24.2 ± 2.3, KD_dev_ 27 ± 2.6; cinnamon: Scramble 27.4 ± 2.3, KD_dev_ 27.8 ± 2.1).

We also performed control experiments to assess the sensory or motor confounds to the observed differences in pre-attentive (Figures S4D-G) and attentional processing (Figures 4E-G, S-T). As with CC mice, KD_dev_ and controls displayed no significant differences in gross motor control, motor coordination, or hearing (Figures 4E-G), nor did they display broad cognitive deficits (Figures 4S-T). Notably, however, in contrast to CC mice, they exhibited no significant differences in anxiety-like behavior (Figure 4U-V). Altogether, these results demonstrate that reducing the expression of *Homer1a/Ania3* in PFC during early postnatal development is sufficient to improve pre-attentive processing and attentional performance in adult. This raises two questions: 1) how does endogenous differential expression of short *Homer1/Ania3* isoforms throughout development affect cellular functions underlying attention in the adult, and 2) how do these cellular and molecular changes influence neural dynamics supporting attention?

### Low Homer1a expressing neurons up-regulate GABA-receptors

To better understand the differences in transcriptional programs associated with *Homer1* we performed single cell RNAseq from PFC of adult CC083 and CC025 mice (Figure 5A). After applying quality control filters (Methods) we obtained 70,920 total cells (Figure S5A; 40,897 from CC083 and 30,023 from CC025, n=2 biological replicates per CC line of 3 mice pooled per replicate). We performed graph-based weighted nearest neighbors clustering analysis and identified major cell types based on cluster-wide expression of several canonical marker genes (Figures 5B and S5B-C)^48^.

**Figure 5.**
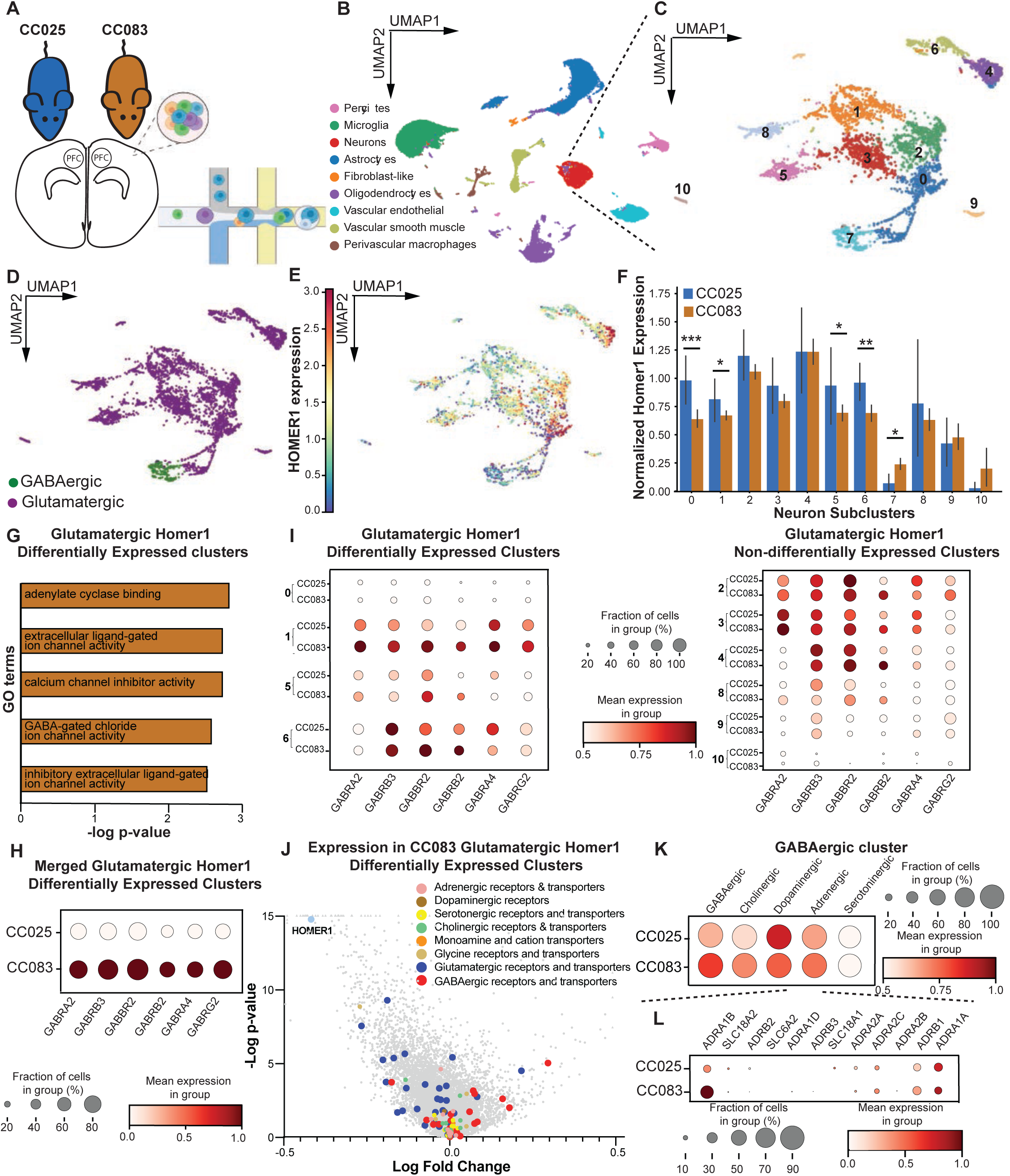
Low Homer1-expressing glutamatergic neurons upregulate GABA receptors. (**A**) Schematic representation of scRNAseq workflow. (**B**) UMAP visualization of all cells collected from CC025 (n=6) and CC083 (n=6) mice clustered based on transcriptional profile. (**C**) UMAP visualization sub-clustering all cells identified as neurons. (**D**) Identification of excitatory (glutamatergic) and inhibitory (GABAergic) neuron clusters based on expression of canonical marker genes. (**E**) UMAP visualization of scaled *Homer1* expression in neuronal clusters. (**F**) Differential *Homer1* expression between CC083 and CC025 neurons by cluster (*p≤0.1, **p≤0.01, ***p≤0.001). Data shown as mean ± SD. (**G**) Gene ontology (GO) analysis of molecular function for genes upregulated in the glutamatergic *Homer1* DE clusters from CC083 mice. (**H-I**) Dot plots showing scaled expression of GABAergic receptors driving GO analysis (from **G**) in (**H**) the glutamatergic *Homer1* DE clusters by line and (**I**) all glutamatergic clusters by cluster and by line. (**J**) Volcano plot depicting differential gene expression in the glutamatergic *Homer1* DE clusters in CC025 and CC083, relative to the CC025. Colored dots indicate genes encoding receptors and transporters of common neurotransmitter systems. (**K**) Dot plot showing the expression of markers for common neuromodulatory systems in GABAergic cluster 7 by line. (**L**) Dot plot of adrenergic receptors and transporters in CC025 and CC083 cells in GABAergic cluster 7. For all dot plots, the size of each dot corresponds to the percentage of cells from each line expressing each gene or gene set, and the color intensity indicates the relative, scaled expression of the gene/gene set.

Since *Homer1* is primarily expressed in neurons^49^, we sub-clustered the neurons (4,633 cells) and re-clustered them based on the first 50 principal components, identifying 10 distinct neuronal clusters (Figure 5C; Methods). We determined that 9 of the clusters were glutamatergic and 1 was GABAergic based on the expression of marker genes *Slc17a6*, *Slc17a7*, *Slc32a1*, and *Gad1* (Figure 5D). Consistent with previous studies^50,51^, *Homer1* expression was primarily restricted to glutamatergic neurons (Figure 5E). Of the 9 glutamatergic clusters, 4 showed substantial downregulation of *Homer1* in CC083 cells compared to CC025 cells (Figure 5F; clusters 0,1,5, & 6 referred to as *Homer1* Differentially Expressed (DE) clusters). To define the gene expression patterns associated with varying levels of *Homer1* we performed differential expression analysis on the *Homer1* DE clusters between CC lines^52^. Gene ontology analysis of molecular function for genes upregulated in the CC083 cells from the *Homer1* DE clusters showed an enrichment of pathways related to inhibitory GABA signaling, while CC025 terms overrepresented glutamatergic signaling, driven by GABA and glutamate receptor subunits, respectively (Figures 5G and S5D). In fact, CC083 cells uniformly upregulate several GABA receptor sub-types (Figure 5H), specifically in the *Homer1* DE clusters (Figures 5I), further apparent when analyzing the Homer1+ cells within those cluster (Figure S5E-F), and meanwhile downregulating several glutamatergic receptor subtypes with almost no differential expression of other neurotransmitter receptors or transporters (Figure 5J). These data indicate that lower expression of *Homer1* in a subset of prefrontal excitatory neurons yields enhanced GABAergic to glutamatergic receptor balance in those same neurons, suggesting enhanced inhibitory receptivity.

We next assessed the transcriptional programs of upstream GABAergic neurons. To do so, we performed differential expression analysis on the GABAergic cluster, in which, interestingly, *Homer1* is significantly upregulated in the CC083s (Figure 5F, p=0.02). Due to the well-studied contributions of neuromodulation in attentional processing^15^, we assessed expression differences of markers for the most common neuromodulatory systems and found that CC083 GABAergic neurons had higher expression of genes associated with adrenergic and cholinergic signaling than the CC025s (Figure 5K). Furthermore, pathway enrichment analysis^53–55^ indicated significant overrepresentation of genes related to noradrenergic signaling in CC083s (Figure S5G). Given its historical significance in attentional regulation^7,12,56,57^, as well as its role as a target of medications to treat ADHD^58,59^, we further analyzed the expression of adrenergic receptors. We found that the higher expression of adrenergic marker genes in CC083 GABAergic cells is driven primarily by the adrenergic receptor *Adra1b*, which, along with *Adrb1*, appear to be preferentially expressed in the GABAergic cluster (Figure 5L).

To determine whether these differences between strains was causally driven by developmental changes in *Homer1a/Ania3* expression, we prepared another cohort of mice with bilateral injection of either *Homer1a/Ania3* shRNA or scrambled controls at P0. We then performed scSeq from adult mice and performed similar sets of analyses as with the CC mice. We found that within the one main cluster of excitatory neurons (Figure S5H-I), *Homer1* was significantly downregulated in cells from the KD_dev_ mice (Figure S5J), while indeed in those same cells the GABA receptor genes were significantly upregulated (Figure S5K). Furthermore, within the inhibitory neuron cluster, KD_dev_ mice displayed strong enrichment of adrenergic receptor subtypes. Taken together, these data demonstrate that developmental prefrontal *Homer1/Ania3* knockdown is causally related to increased prefrontal receptivity to adrenergic and inhibitory input. We next explored the consequences of these effects on neural dynamics in the behaving animal during an attention task.

### Developmental reduction of *Homer1/Ania3* alters prefrontal inhibitory influence, enhances SNR, and improves attention

How do the *Homer1a*-associated molecular and cellular changes contribute to changes in neural dynamics underlying attentional processing? Are there roles for long-range inhibitory recurrence via thalamus, or neuromodulation via locus coeruleus, or both, linked to attentional performance? And are there contributions of non-neuronal cells (i.e., oligodendrocyte precursor cells (OPCs) or mature oligodendrocytes (OLs) (Figures S6A-D) in regulating prefrontal dynamics and attentional processing? To address these and other questions, we moved to an *in vivo* preparation to record multi-area brain activity as CC083 (low *Homer1a*, high attention) and CC025 mice (high *Homer1a*, low attention) performed the operant signal detection task. We injected AAV1/9-GCaMP or JRGECO1a into locus coeruleus (LC, GCaMP), mediodorsal thalamus (MD, GCaMP), PFC oligodendrocytes (OL, GCaMP; validation of OL-specific AAV-MAG-GCaMP in Figures S6E-F) and PFC neurons (PFC, JRGECO1a), implanted optical fibers above each region, and used a custom dual-color fiber photometry system to record bulk calcium signals from these regions simultaneously in behaving mice (Figure 6A). Because CC025 mice did not tolerate intracranial implants we used B6 mice as “low performers” since they have comparable *Homer1a* expression and behavioral performance to CC025s (Figures S6G, 1B, and 2E). Multi-area neural activity recordings from a given animal were frame projected onto a camera sensor, and custom scripts (Methods) were used to extract time-series data, regress out motion-related artifacts, and align to behavioral data (example alignment from one trial of a CC083 mouse in Figure 6B).

**Figure 6.**
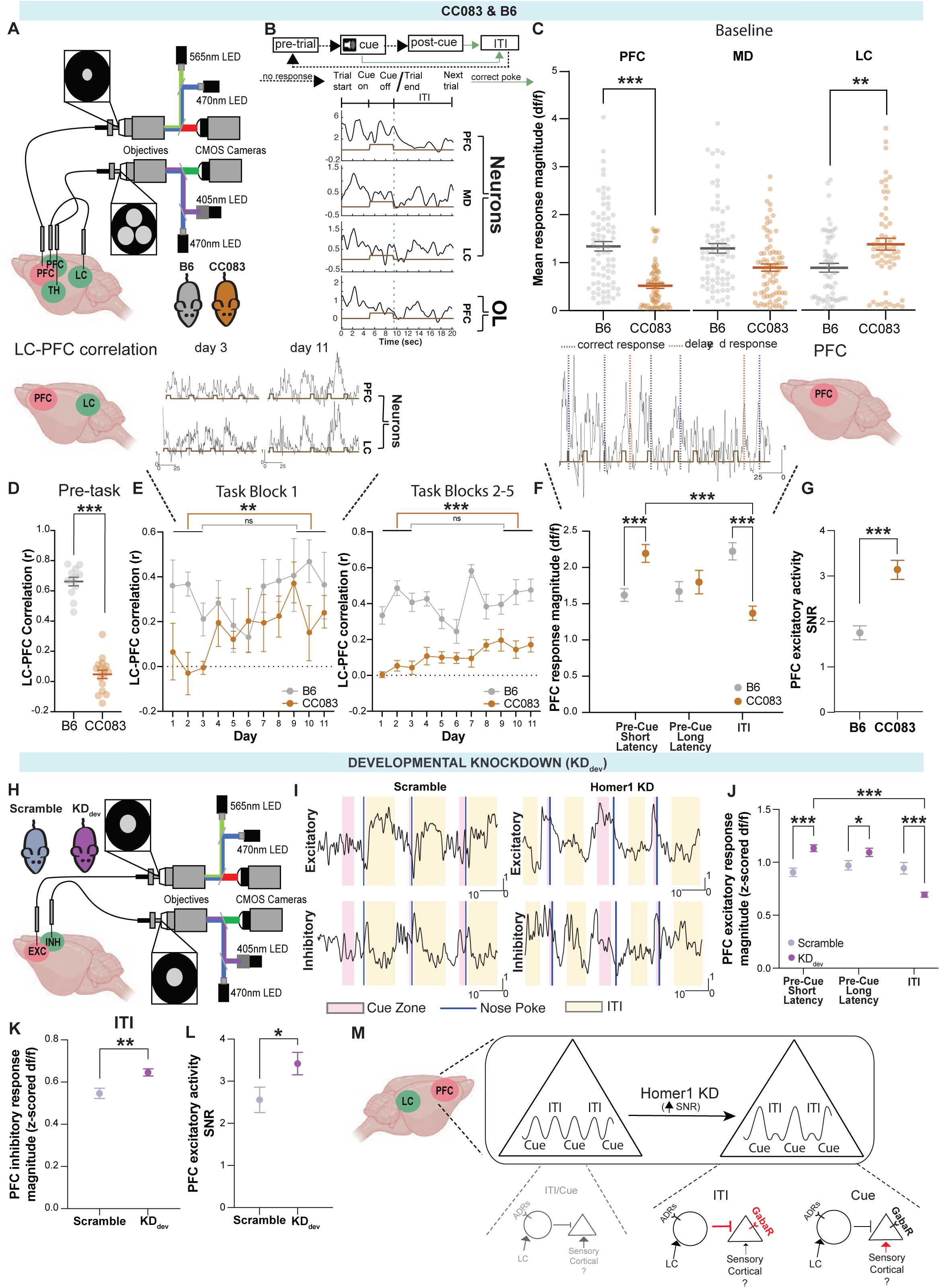
Developmental reduction of *Homer1/Ania3* alters prefrontal inhibitory influence, enhances SNR, and improves attention. (**A**) Schematic of dual-color, 4-region photometry system. Simultaneous 565 nm, 470 nm, and 405 nm recordings were taken from PFC neurons (jRGECO, red), MD neurons, LC neurons, and PFC oligodendrocytes (GCaMP, green) in B6 (grey) or CC083 (tan) strains. (**B**) Top: Schematic representation of SDT trial structure. Bottom: example jRGECO (PFC neurons, row 1) and GCaMP (MD neurons, LC neurons, and PFC oligodendrocytes (OL; rows 2-4, respectively) traces aligned to a SDT trial. (**C**) Average activity during baseline for B6 (n=4) and CC083 (n=4 for PFC and MD, n=3 for LC) during 1 min recordings (Welch-corrected t-test, p(PFC)<0.001, p(MD)=0.002, p(LC)=0.002). (**D**) Pairwise Pearson’s correlations between LC and PFC neuronal activity at baseline (Welch-corrected t-test for B6 *vs* CC083, n=4 each, 5 min recordings, p<0.001). (**E**) Top: representative traces from PFC (top) and LC neurons (bottom) from day 3 (left) and 11 (right), Y-axis is z-scored df/f and X-axis is time (s). Brown rectangles indicate cues. Bottom: Pairwise Pearson’s correlations between LC and PFC activity during SDT sessions. Each 20 min session was split into 5, 4-min blocks. Data is shown from the first 4-minute block (left) and for blocks 2-5 (right) as mean ± SEM (Welch-corrected t-tests for days 1-3 *vs* days 9-11 within strain, for CC083 p(block1)=0.003 and p(blocks 2-5)<0.001). (**F**) Top: Representative trace from PFC, Y-axis is z-scored df/f and X-axis is time (s). Brown rectangles indicate cues, orange dotted lines indicate delayed responses, blue dotted lines indicate correct responses. Bottom: PFC activity in task during the 5 sec before cue onset of short and long latency response trials, respectively, and during the last 5 seconds of ITIs for trials on all days in B6 (n=5) and CC083 (n=4) mice (two-way ANOVA, followed by Holm-Sidak’s test for multiple comparisons, *** indicates p<0.001). (**G**) PFC neuronal signal to noise (SNR: trial pre-cue maximum − mean-baseline) / SD-baseline) 5 seconds before cue onset in B6 (n=5) and CC083 (n=4) mice for correct trials on all days (Welch-corrected t-test, p<0.001). (**H**) Schematic of dual-color recordings from PFC excitatory neurons (jRGECO, red) and inhibitory neurons (GCaMP, green) in Scramble and KD_dev_ mice. (**I**) Example traces from excitatory (top) and inhibitory (bottom) neurons across 3 trials in Scramble (left) and KD_dev_ (right) mice. (**J**) PFC excitatory activity in task during the 5 sec before cue onset of short and long latency response trials, respectively, and during the last 5 seconds of ITIs for trials on all days in Scramble (n=6) and KD_dev_ (n=10) mice (two-way ANOVA, followed by Holm-Sidak’s test, *** indicates p<0.001, * indicates p=0.03). (**K**) PFC inhibitory activity in task during the last 5 seconds of ITIs for trials on all days in Scramble (n=6) and KD_dev_ (n=10) mice (Welch-corrected t-test, p=0.001). (**L**) PFC excitatory neuronal SNR 5 seconds before cue onset in Scramble (n=6) and KD_dev_ (n=10) mice for correct trials on all days (Welch-corrected t-test, p=0.04). (**M**) Putative model: Knockdown of Homer1 improves SNR by reducing PFC activity during baseline periods of a task (here depicted as inter-trial intervals, ITIs) but dynamically elevating activity during cue-presentations. Reduced *Homer1/Ania3* levels leads to increased GABA receptor expression in excitatory neurons, and increased inhibitory tone in PFC (either by increasing feed-forward inhibition from LC, or how the excitatory neurons sense ongoing inhibition, or both) during non-attentive baseline periods of a task. When attention is required, incoming excitatory input overrides ongoing inhibition to provide targeted cue-related responses.

We first analyzed baseline activity patterns in CC083 and B6 mice and noticed substantially depressed PFC activity (p<0.001, Welch-corrected t-test) and elevated LC activity (p=0.002, Welch-corrected t-test) in CC083 mice (Figure 6C). This, together with the observation from scSeq data of increased adrenergic *Adra1b* reception onto PFC GABAergic cells (Figure 5L), suggested that LC may contribute to baseline inhibitory tone in PFC through feed-forward inhibition. Indeed, we found that LC-PFC functional correlations when not engaging in a task were close to Pearson’s r=0 in CC083s compared to ~0.7 in B6 (Figure 6D, p<0.001, Welch-corrected t-test). However, as mice started training on the signal detection task, we noticed a steady increase in LC-PFC functional correlations in CC083 mice (Figure 6E) that mirrored their steady improvement in task performance (Figure S6H), and which were unlikely to be due to contamination of LC fibers in PFC (since these recordings were made on contralateral sides, using separate sensors and imaging filters, and emerged only in certain phases of the task, Methods). This steady increase in LC-PFC correlations were not observed in B6 mice, presumably due to already high baseline correlations precluding further enhancements during task (Figure 6E-F, multiple Welch-corrected t-tests p(block 1)=0.009, p(blocks 2-5)<0.001; example raw traces from day 3 and day 11 are shown). This improved LC-PFC coupling in CC083 mice was strongest in the first four minutes (block 1) of each day’s session (Figure 6E, left panel), after which these correlations reduced back toward baseline (blocks 2-5; Figure 6E, right panel).

In addition to increased LC-PFC coupling, we found that during the task CC083s exhibited large increases in PFC activity before and at cue-onset, which were greater on shorter latency correct trials compared to long latency trials and omissions (Figure 6F, example raw traces from correct and incorrect trials shown). More strikingly, this cue-related activity rapidly diminished during inter-trial intervals (ITIs) (Figure 6F). Such dynamic task related fluctuations in CC083s led to consistently high levels of PFC signal to noise (SNR: trial averaged neural activity in cue vs baseline periods of the task) throughout the task (Figure 6G). Notably, these dynamic task-related fluctuations in PFC activity, and enhanced SNR, were not observed in B6 mice, which exhibited relatively constant PFC responses throughout all task phases, including inter-trial intervals, short and long latency responses (Figures 6C, 6F, and 6G).

To test whether these dynamic task related fluctuations and enhanced SNR in CC083 mice were causally driven by changes in Homer1, we prepared a new cohort of B6 mice for photometry with developmental knockdown of prefrontal *Homer1a/Ania3* compared with Scramble controls. We simultaneously recorded excitatory neurons in PFC using jRGECO and inhibitory neurons in PFC using mDlx-GCaMP as mice performed the operant SDT (Figures 6H-L, S6I-N). The results from these experiments beautifully recaptured the SNR effects that we had observed in CC083 vs B6 mice (Fig. 6F-G and 6J-L). Specifically, PFC excitatory responses were substantially higher at cue-presentation than during ITI, specifically in KD_dev_ mice compared with controls (Fig. 6I,J), leading to significant improvements in SNR during task performance (Fig. 6L). One notable difference between the CC083s and KD_dev_ mice is that the baseline inhibitory tone in KD_dev_ mice was reflected acutely during the task (during ITIs) whereas more chronic inhibitory tone was observed in the CC083s, which were apparent even outside of the task and during home-cage (Figure 6C vs Figure S6K). This may reflect acute compensation of Homer1 knockdown in KDdev mice or that other effects beyond the locus and gene contribute to a more chronic inhibitory tone in CC083s, which is also reflected in their even higher attentional performance (changes in latency as well as omissions leading to impressive overall increases in task accuracy; Figures S6M-N).

Focusing next on the inhibitory neurons, we found a small but significant increase in the activity of inhibitory neurons during ITIs in KD_dev_ mice compared with controls (Fig. 6K), though these were not as striking as the large magnitude changes in excitatory responses during task (Fig. 6J, short latency cue vs ITI). Together, these data suggest that *Homer1a/Ania3* improve prefrontal SNR by dynamically scaling excitatory neuron activity (low during ITIs but high at cue-presentation), which is in part facilitated by an increase in inhibitory activity (Figures 6I and K), but also a greater sensitivity of excitatory neurons to this inhibition (Figures 6J and L, also supported by striking elevations of GABARs specifically in KD_dev_ cells, Figures S5K).

Taken together, these results suggest a model in which low *Homer1a*/*Ania3* enhance inhibitory receptivity, allowing for dynamic scaling of prefrontal activity, and targeted elevations at cue-onset, linked to short-latency correct responses (Figure 6M). These results therefore reveal that the critical component of prefrontal processing during attention may not be overall increases in activity, but rather frequent cue vs ITI resets that enable increased SNR and targeted behavioral responses.

## DISCUSSION

Here we performed genetic mapping in outbred mice and identified a short segment on chromosome 13 that explains a large portion of the variance on a pre-attentive processing task. Within this locus, we identified a causative gene as Homer1, synaptic protein with known roles in regulating excitatory glutamatergic transmission. In particular, knockdown of the activity-dependent short-isoforms of Homer1 – *Homer1a/Ania3* – in prefrontal cortex during a developmental period led to significant improvements in pre-attentive processing and multiple measures of attention in the adult. Notably, the effects of *Homer1a/Ania3* were highly specific to attention, as there were no overall changes in the ability to learn the tasks, and perform other cognitive functions, nor were there obvious sensory-motor impairments or changes in measures of anxiety. We believe the specificity of these behavioral effects on attention are due to the isoform, region-, and developmental-window specific perturbations of *Homer1/Ania3*. How variation at the chromosome 13 locus leads to such targeted cell-type and isoform specific changes in gene-expression of an otherwise ubiquitously expressed gene will be an important avenue of future investigation.

A rich history of work on Homer1 and its isoforms have revealed important roles in excitatory neurotransmission affecting multiple cognitive domains^43–46^, but little is known about its role in attention, particularly by sculpting inhibition and during a defined developmental window. Notably, genes related to Homer1 signaling have been repeatedly identified in human studies linked to ADHD, suggesting a core underlying biology for attention^60–63^. In addition to ADHD, *Homer1a* is also associated with schizophrenia^64,65^ and autism^66,67^, suggesting that early dysfunctions in sensory gating (leading to sensory overload) provide common etiology driving diverse downstream neuropsychiatric symptoms characterized by hallucinations, hypersensitivity, and motor compensations. Thus, prefrontal Homer1 may be a hub for deeper mechanistic understanding of sensory gating and attention, indeed it has outsized contribution to the trait and may therefore point us toward unifying circuit models.

In our initial attempts to understand how *Homer1a* might shape behavioral improvements in attentional performance, we explored the molecular programs associated with *Homer1a* expressing neurons as well as their associated circuit physiology in the context of their inputs and outputs. Through cellular resolution RNA sequencing analysis, we found that low-*Homer1a,* high attention mice (CC083) only down-regulate *Homer1* expression in a subset of PFC excitatory neurons, which in turn is associated with significant upregulation of GABAergic receptors in these same cells. Moreover, knocking down *Homer1a* locally in PFC during postnatal development led causally to similar changes in GABAergic receptor upregulation. Assuming these receptors receive inputs from local GABAergic neurons, we further analyzed these inhibitory neurons and found enrichment of a specific adrenergic receptor gene, *Adra1b*, in these cells. These data together suggested that in mice with high attentional performance, chronic downregulation of *Homer1a* may drive homeostatic scaling^68–71^, favoring inhibitory inputs and overall inhibitory tone in PFC.

Indeed *in vivo* neural activity recordings revealed strongly reduced baseline PFC activity, which was present chronically in high attention CC083 mice, and more acutely during baseline periods of task (ITIs) in mice with developmental knockdown of Homer1a. However, during attentionally-demanding periods of task performance, this inhibitory influence was relieved and large increase in PFC cue-related activity enabled short-latency correct responses. Notably high-*Homer1a*, low-attentional CC performers, exhibited uniformly elevated PFC activity at both baseline and cue related phases of the task. Thus, rather than overall increases in PFC neural activity, a dynamic prefrontal inhibitory influence, increased SNR, and targeted cue-specific response enabled attentional performance. Given that widely prescribed medications for ADHD are stimulants acting to elevate PFC activity, which, while effective, can lead to rapid tolerance, a strategy to reduce PFC activity and tune its SNR may be therapeutically promising.

While the current investigations reveal mechanisms of attention related to the interplay of prefrontal inhibitory tone on increased SNR of PFC, an understanding of the more complete effects of *Homer1a* require deeper investigation. For instance, GABAergic cells from low-*Homer1a* mice also upregulate cholinergic signaling suggesting cross-talk between neuromodulatory systems, likely over diverse time-scales. We also note that throughout the study there were interesting links between oligodendrocyte function and attentional performance that may be important to investigate further. For instance, the top 20 genes upregulated in PFC of the DO high performers mapped almost exclusively to expression in the oligodendrocyte lineage (Figures S6A-B), and a similar shift in oligodendrocyte transcriptional programs was also noticeable between the CC high and low performers (Figures S6C-D). Furthermore, preliminary recordings in PFC oligodendrocytes (Figures S6E-F) revealed dynamic task-dependent changes in calcium responses, supporting roles for activity-dependent adaptive myelination in attentional processing^72–75^. In sum, the identification of a single gene with large contributions to attention, and its roles in tuning prefrontal inhibitory tone and SNR, provides tractable inroads into circuit models and non-stimulant based therapeutic strategies for attentional processing.

### Limitations of the study

To facilitate the simultaneous recording of multiple cell types and brain areas during behavior we opted to perform bulk photometry-based neural recordings rather than cellular resolution imaging. This approach, while enabling cross region and cell-type analyses, prevented a more highly resolved analysis of the role of single cells within the prefrontal microcircuit, which we aim will be the focus of future studies. Additionally, it is important to acknowledge that strain-specific differences may be contributing to the observed difference in dynamic range of the LC-PFC functional correlations. Such strain-based differences would be important to investigate further. Finally, due to technical and practical considerations the potential adrenergic contribution to our model is based on both single-cell sequencing data and LC recordings, but simultaneous, direct measurements of local norepinephrine release would be ideal for future studies to provide a more accurate representation of neurotransmitter dynamics.

## ACKNOWLEDGMENTS

We thank Cori Bargmann, Jeff Friedman, and Rick Lifton for helpful discussions related to genetics and genetic-mapping in mice. We thank Jim Hudspeth for advice on testing auditory brainstem responses, and Mary Beth Hatten and Bob Darnell for discussions related to developmental Homer1 functions. We thank Josue Regalado for help with photometry experiments and analysis, Jazz Weisman for contributions to the dual-color photometry system, and Vivian Li for stereotactic surgeries. We thank Paul Greengard’s laboratory for sharing behavioral instrumentations and Roshan Sharma for suggestions on the scRNAseq analysis. We are thankful to core facilities at Rockefeller (Precision Instrumentation, Genomics, & FACS), MSKCC (Single Cell Analytics Innovation Lab), and University of Arizona (Viral Production Core). Cartoons in Figures 3, 4, 6, and S3 were created with biorender.com. This work was supported by a Kavli Institute pilot grant from the Rockefeller University (AT), a Medical Scientist Training Program grant from NIGMS of the NIH (T32GM007739) to the Weill Cornell/Rockefeller/Sloan Kettering Tri-Institutional MD-PhD Program. (AFI), and grants from the National Institutes of Health under award numbers 5R01MH110553 (NVDM), DP2AG058487 (PR), and from the Harold & Leila Mathers Foundation (PR).

## AUTHOR CONTRIBUTIONS

ZG, PR, and PS conceived the study. ZG, ABO, and PR designed the experiments. ZG selected and optimized the behaviors used in this study, performed mouse behaviors (together with ABO), molecular studies including RNA preparation and single cell dissociation (together with AT), cloning and cell-based assays, and *in vivo* neural activity recordings and analysis, supervised by PR. MK performed QTL analysis as well as RNAseq and scRNAseq analysis, supervised by PS. AT assisted with designing and performing the scRNAseq experiments and analysis. ZG and GR designed the head-fixed behaviors and GR performed these experiments. JF, YH, and MG assisted with surgeries and histology. AFI performed surgeries for the developmental study, supervised by ND. BF performed auditory brainstem recordings, supervised by AJH. ZG and PR wrote the paper with input from all authors.

## STAR METHODS

### Key Resources Table

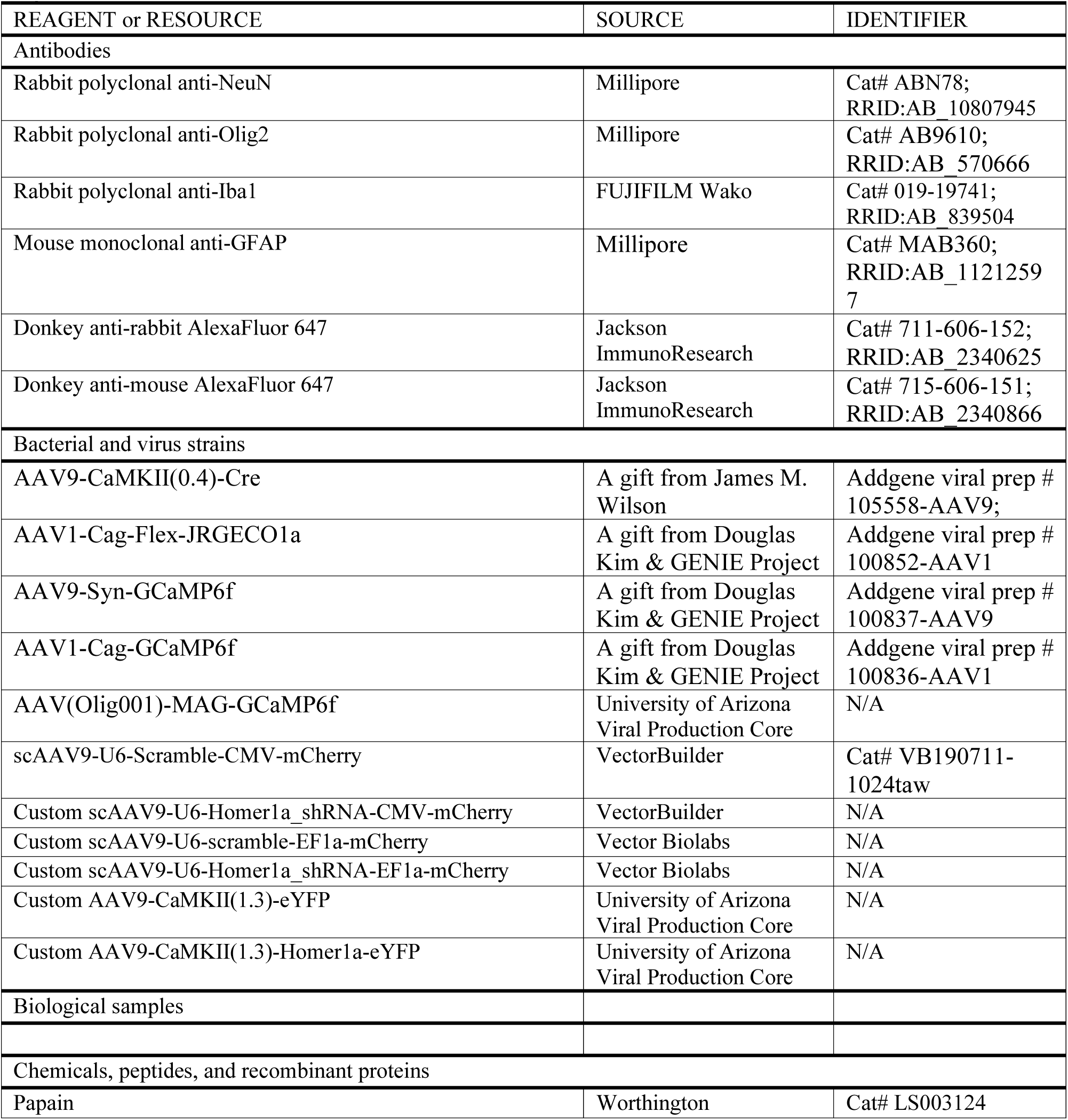
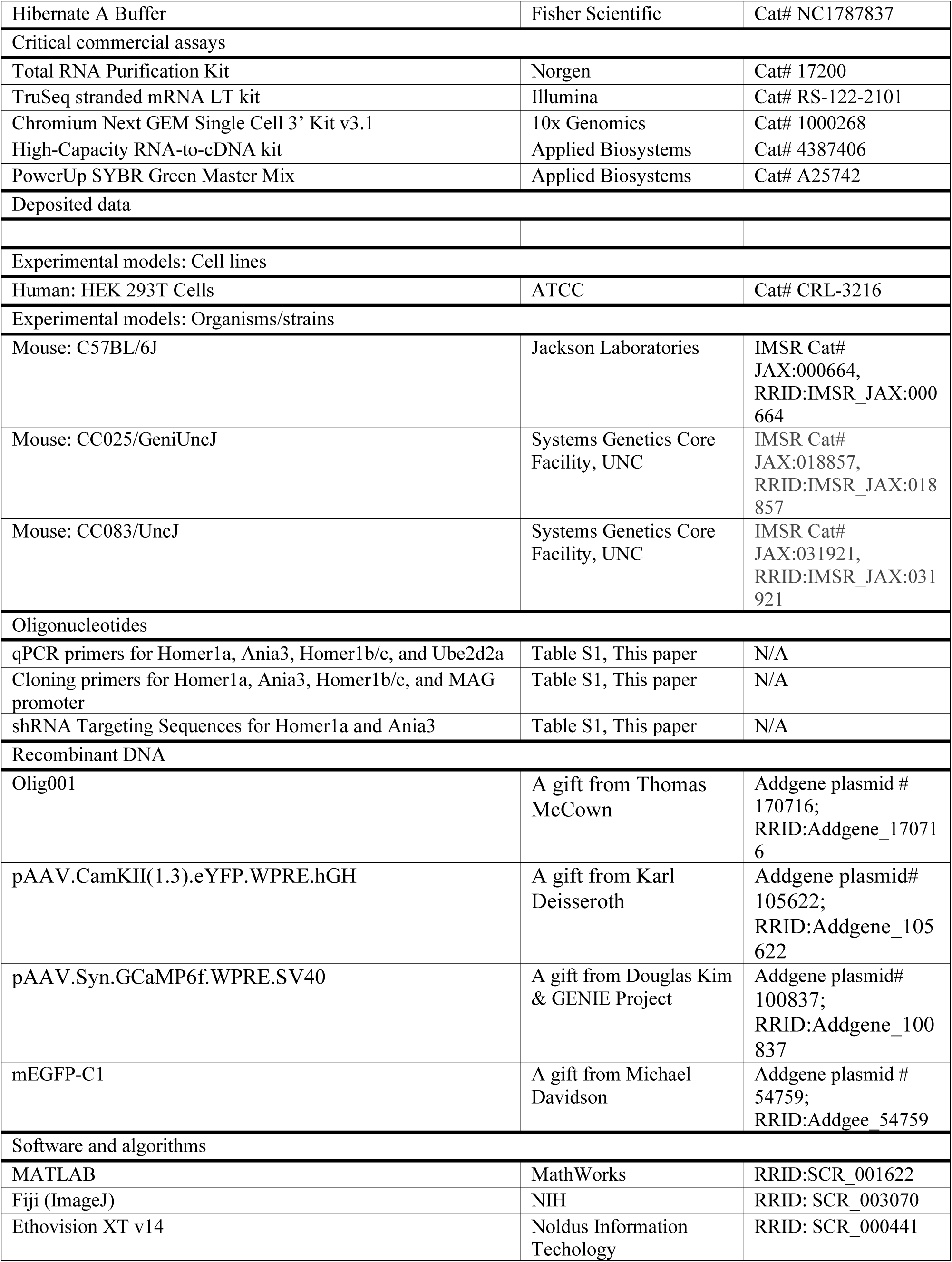
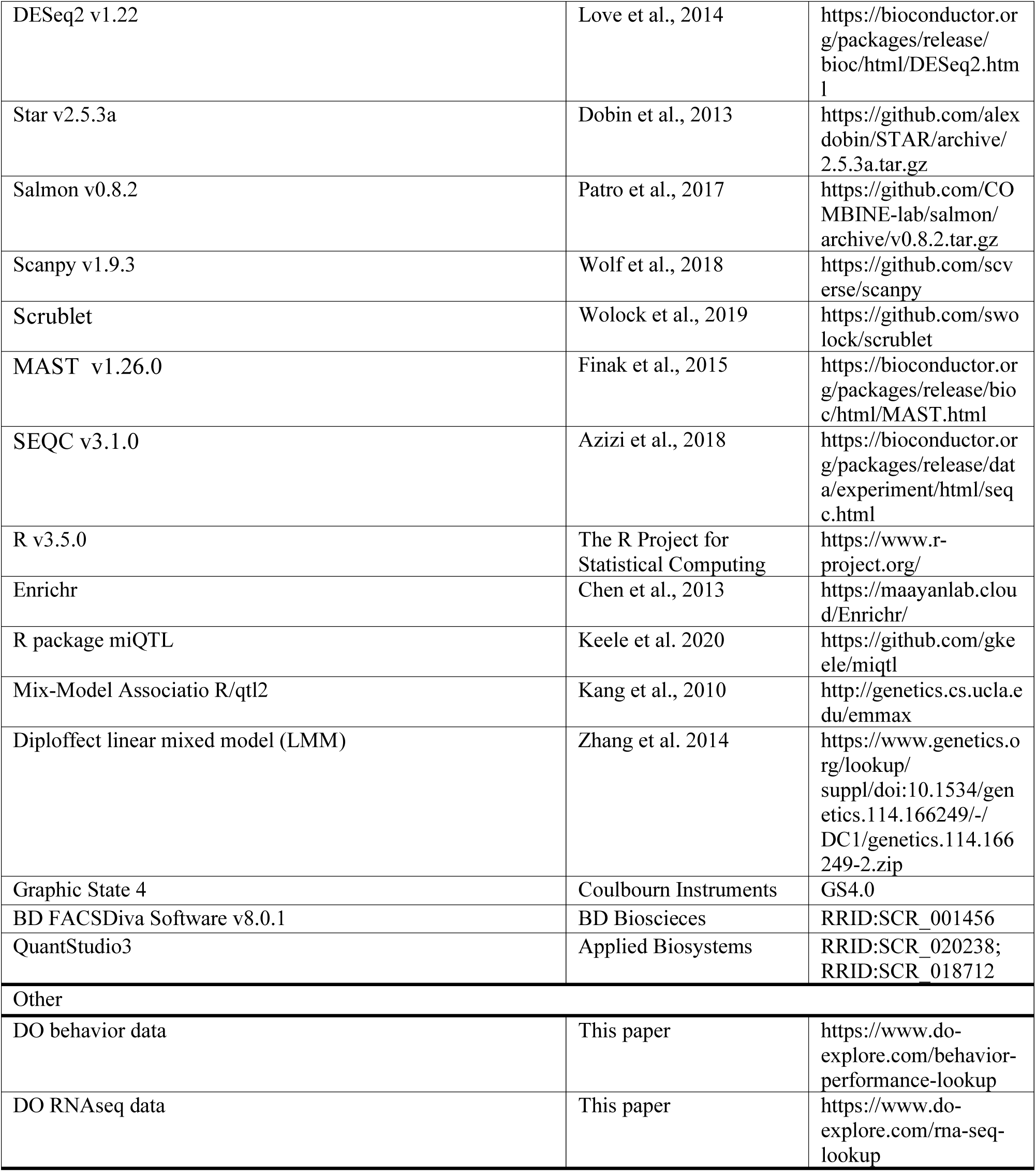

## RESOURCE AVAILABILITY

### Lead Contact

Further information and requests for resources and reagents should be directed to and will be fulfilled by the Lead Contact, Priyamvada Rajasethupathy.

### Materials Availability

All other unique reagents generated in this study are available from the Lead Contact with a completed Materials Transfer Agreement.

### Data and Code Availability

Numerical data for each figure are included with the manuscript as source data. All other data are available from the corresponding author upon request. Custom MATLAB code are available upon request from the Lead Contact.

## EXPERIMETAL MODEL AND SUBJECT DETAILS

### Animals

C57Bl6/J (B6) and Diversity Outbred (DO, 25th generation) male mice were purchased from The Jackson Laboratory. Collaborative Cross (CC) male mice from the CC083 and CC025 lines were purchased from the University of North Carolina at Chapel Hill. All mice were bought at six to eight weeks old, group housed three to five per cage and kept under a 12 h light–dark cycle in a temperature-controlled environment with ad libitum food and water, unless mice were food restricted for behavioral assays. All procedures were done in accordance with guidelines approved by the Institutional Animal Care and Use Committees (protocol #22087-H) at The Rockefeller University. Number of mice used for each experiment was determined based on expected variance and effect size from previous studies and no statistical method was used to predetermine sample size. DO phenotyping was performed with all males to sufficiently power the study at affordable cost, but future studies will use female-only or mixed cohorts.

## METHOD DETAILS

### Surgical Procedures

Surgical procedures and viral injections were carried out under protocols approved by Rockefeller University IACUC and were performed in mice anesthetized with 1%–2% isoflurane using a stereotactic apparatus (Kopf) under a heating pad. Paralube vet ointment was applied on the eyes to prevent drying.

#### Viral Injections

Virus was injected using a 34–35 G beveled needle in a 10ul NanoFil Sub-Microliter Injection syringe (World Precision Instruments) controlled by an injection pump (Harvard Apparatus). All viruses were injected at a volume of 1 µL and a rate of 100nL/min (unless otherwise mentioned), and the needle was removed 10 min after the injection was done to prevent backflow of the virus. All injection coordinates were relative to bregma.

For adult knockdown manipulations: B6 mice were bilaterally injected at the age of 8 weeks in the PFC (A/P: 1.8 mm, M/L: ±0.3 mm, D/V: −1.75 mm) with an scAAV9 expressing either a U6-scramble (non-targeting) shRNA-CMV-mCherry (titer: 9.87. × 10^12^ GC/mL, VectorBuilder) or U6-Homer1a-targeted shRNA-CMV-mCherry (titer: 4.8 × 10^12^ GC/mL, VectorBuilder) construct.

For adult overexpression manipulations: B6 mice were bilaterally injected (2 injections/hemisphere) at the age of 8 weeks in the PFC (A/P: 1.8 mm, M/L1: ±0.3 mm, M/L2: ±0.45 mm, D/V: −1.75 mm) with an AAV9 expressing either CaMKII(1.3)-eYFP (titer: 1.0 × 10^13^ GC/mL) or CaMKII(1.3)-Homer1a-eYFP (titer: 1.0 × 10^13^ GC/mL) construct at a volume 0.5 µL for each injection. pAAV.CamKII(1.3).eYFP.WPRE.hGH was a gift from Karl Deisseroth (Addgene plasmid# 105622; http://n2t.net/addgene:105622; RRID:Addgene_105622).

For developmental knockdown experiments: Injections in pups were performed according to previously described anesthesia and injection protocols^76^. Here, B6 pups were bilaterally injected in PFC at p0 (A/P: ~0.3 mm, M/L: ~±0.15-0.2 mm, ~-0.7-0.8 mm) and again at p11 (A/P: 0.51, M/L: ±0.17, D/V: −1.5 mm) with an AAV9 expressing either a U6-scramble (non-targeting) shRNA-EF1a-mCherry (titer: 4.8. × 10^12^ GC/mL) or pooled U6-Homer1a-targeted shRNA-EF1a-mCherry and U6-Ania3-targeted shRNA-EF1a-mCherry construct (titer: 2.8 × 10^12^ GC/mL, Vector Biolabs) construct at a volume of 0.1 µL both times.

For multi-fiber photometry experiments: A mixture of AAV9-CaMKII(0.4)-Cre (titer: 1.0 × 10^13^) and AAV1-Cag-Flex-JRGECO1a (titer: 1.0 × 10^13^) was injected into PFC (A/P: 1.85 mm, M/L: 0.35 mm, D/V: −2.55 mm) at a combined volume of 1 µL. AAV9-Syn-GCaMP6f (titer: 1.4 × 10^13^ GC/mL) was injected ipsilaterally into MD (A/P: −1.6 mm, M/L: 0.45 mm, D/V: −3.2 mm). AAV(Olig001)-MAG-GCaMP6f (titer: 1 × 10^13^ GC/mL, Univ. Arizona Viral Production Core) was injected into PFC contralaterally (coordinates: A/P: 1.85 mm, M/L: −0.35 mm, D/V: −2.55 mm). AAV1-Cag-GCaMP6f (titer: 2.6 × 10^12^) was also injected contralaterally to the initial injection (Cag-Flex-JRGECO1a) into LC (A/P: −5.4 mm, M/L: −0.85 mm, D/V: −3.6 mm). pENN.AAV.CamKII 0.4.Cre.SV40 was a gift from James M. Wilson (Addgene viral prep # 105558-AAV9; http://n2t.net/addgene:105558; RRID:Addgene_105558), pAAV.CAG.Flex.NES-jRGECO1a.WPRE.SV40 was a gift from Douglas Kim & GENIE Project (Addgene viral prep # 100852-AAV1; http://n2t.net/addgene:100852; RRID:Addgene_100852;)^77^, pAAV.Syn.GCaMP6f.WPRE.SV40 was a gift from Douglas Kim & GENIE Project (Addgene viral prep # 100837-AAV9; http://n2t.net/addgene:100837; RRID:Addgene_100837), pAAV.CAG.GCaMP6f.WPRE.SV40 was a gift from Douglas Kim & GENIE Project (Addgene viral prep # 100836-AAV1; http://n2t.net/addgene:100836; RRID:Addgene_100836)^78^, Olig001 was a gift from Thomas McCown (Addgene plasmid # 170716; http://n2t.net/addgene:170716; RRID:Addgene_170716)^79^.

#### Cannula implants

Immediately following viral injections, mice undergoing photometry experiments were implanted with 1.25 mm ferrule-coupled optical fibers (0.48 NA, 400 μm diameter, Doric Lenses) cut to the desired length so that the implantation site is ~0.2 mm dorsal to the injection site. Cannula implants were slowly lowered using a stereotaxic cannula holder (Doric) at a rate of 1 mm/min until it reached the implantation site, 0.2 mm dorsal to the injection site. Optic glue (Edmund Optics) was then used to seal the skull/cannula interface and a custom titanium headplate was glued to the skull using adhesive cement (Metabond).

Mice recovered for 5 weeks after Homer1 manipulations and 3 weeks after photometry implants before experiments began.

### Animal Behaviors

#### Acoustic Startle Response and Prepulse inhibition

Acoustic startle response and prepulse inhibition testing was performed as described previously^80^. Startle was measured using a San Diego Instruments SR-Lab Startle Response System. Mice were placed into Plexiglas cylinders resting on a Plexiglas platform with the chamber light on for the entire duration of the experiment. Acoustic stimuli were produced by speakers placed 33 cm above the cylinders. Piezoelectric accelerometers mounted under the cylinders transduced movements of the mice, which were digitized and stored by an interface and computer assembly. Beginning at startle stimulus onset, 65 consecutive 1 ms readings were recorded to obtain the amplitude of the mouse’s startle response. For the acoustic startle sessions, the intertrial interval between stimulus presentations averaged on 15 s (range: 7–23 s). A 65 dB background was presented continuously throughout the session. Startle pulses were 40 ms in duration, prepulses were 20 ms in duration, and prepulses preceded the pulse by 100 ms (onset– onset). The Plexiglas holders were wiped clean and allowed to dry between runs. The acoustic startle sessions consisted of three blocks. Sessions began with a 5 min acclimation period followed by delivery of five startle pulses (120 dB). This block allowed startle to reach a stable level before specific testing blocks. The next block tested response threshold and included four each of five different acoustic stimulus intensities: 80, 90, 100, 110, and 120 dB (data not shown) presented in a pseudorandom order. The third block consisted of 42 trials including 12 startle pulses (120 dB) and 10 each of 3 different prepulse trials (i.e., 68, 71 and 77 dB preceding a 120 dB pulse). PPI was calculated as follows using the trials in the third block: 100 − ([(average startle of the prepulse + pulse trials)/average startle in the pulse alone trial] × 100). In all experiments, the average startle magnitude over the record window (i.e., 65 ms) was used for all data analysis.

#### T-Maze

Tests consisted of a single 5 min trial, in which the mouse was allowed to explore all three arms of a Y maze (arm dimensions: 12” × 3” × 5” in (L × W × H) for 6 minutes while being recorded using a ceiling mounted camera under red light illumination. Mice were acclimated to the experimental site for 1 hr before all experiments. Whenever possible, the experimenter was blind to the viral condition of all mice during behavioral testing, with the exception of CC083 vs CC025 tests due to the difference in their coat color. The animal behavior was automatically tracked and analyzed by the EthoVision XT (Noldus) software for 1) total number of entries into each arm, 2) sequences of arm entries, 3) and distance moved (inch). Correct alternation (% of total number of arm entries) was defined as consecutive entries in 3 different arms. Total number of entries into each arm as well as total distance moved in the apparatus served as controls to exclude confounding factors to the memory performance, such as arm bias and/or differences in gross motor activity.

#### Rotarod

For this task, on day 1 the mice are habituated to the apparatus by being placed on a rod moving at a constant speed of 4 RPM for 5 min. On day 2, the mice are placed on the rod that this time is moving with an accelerating speed from (4 to 40 RPM), for 4 consecutive trials. In each trial, the latency (s) to fall from the rod is measured by an experimenter. The cut off time is 300 seconds. The latency to fall is averaged across trials and used as a measure of motor coordination.

#### Auditory Brainstem Recording Threshols (ABRs)

The mice were anesthetized with ketamine (110 mg/kg) and xylazine (11 mg/kg) via intraperitoneal injection prior to all procedures. Once a suitable plane of anesthesia was reached, one mL of chilled 0.9% sodium chloride was injected into the mouse’s back for hydration. Both eyes also were moistened with ophthalmic ointment (Puralube®, Dechra Veterinary Products). The anesthetized animal was then placed in a sound-isolated, electrically-shielded box on top of a heating pad (40-90-2-05, FHC, Inc.). In conjunction with the heating pad, a rectal probe and DC temperature controller (41-90-8D, FHC, Inc.) were used to maintain the mouse’s temperature near 38 °C. Needle electrodes (GRD-SAF, The Electrode Store) were subdermally placed behind the pinna of the tested ear (reference electrode), in the scalp between the ears (active electrode), and in the back near the tail (ground electrode). ABRs were evoked by tone bursts of 4, 8, 16, and 32 kHz produced by a closed-field magnetic speaker connected to a power amplifier (MF1 and SA1, Tucker-Davis Technologies). Each 5-ms burst was presented 33.3 times per second with alternating polarity. The onset and offset of each burst was tapered with a squared cosine function. For each frequency, the sound pressure level was lowered from 80 dB SPL in 5–10 dB steps until the threshold was reached. If 80 dB SPL was not enough to elicit a response, higher intensities were produced. The entire sound delivery system was calibrated with a ¼ inch condenser microphone (4939-A-011 and 2690-A-0S1, Brüel & Kjær). The electrical response evoked by the tone bursts and measured by the needle electrodes was amplified 10,000 times and bandpass filtered at 0.3–3 kHz (P55, Astro-Med Inc.). The amplified response was then digitally sampled at 10-μs intervals with a data acquisition device (PCI-6259, National Instruments) controlled by custom software (LabVIEW 2019, National Instruments). The electrical responses to 1,000 bursts were averaged at each intensity level to determine the threshold, which was defined as the lowest level at which any response peak was distinctly and reproducibly present. Visual inspection of the vertically stacked responses facilitated threshold determination.

#### Signal Detection Task (SDT)

Experiments were performed within a Habitest Modular Arena and controlled/recorded by Graphic State 4 software (Colbourn).

Three days prior to the experiment, mice were gradually food restricted to 85% of their body weight by providing ~2g of food per mouse per day and habituated to chocolate pellets by providing 2/3 pellets per mouse per day in their home cage. From the start of food deprivation and for the entire duration of the experiment, body weight and overall well-being were monitored by daily observation and weighting. All training and testing occur immediately before daily feeding.

The protocol is divided into multiple phases:

- ***Magazine Shaping*.** The box is configured to have the chocolate pellet magazine and dispenser, the white LED chamber light, speaker. The mouse enters the box with chamber light off. A reward pellet is dispensed into the magazine on a variable 7-13 sec (VI. 10) schedule and at the same time the magazine light goes on. If the mouse retrieves the pellet, the program returns to VI 10 schedule of reward delivery. Alternatively, if the mouse does not retrieve the pellet within a variable 1-5 min period, the program returns to VI 10 schedule of reward. The session ends after 20 min. When 75% of the cohort are retrieving ≥15 pellets during the magazine shaping phase, the experiment moves to the next phase (usually 1 or 2 days).
- ***Nose Poke Shaping*.** The box configuration is enriched by the nosepoke port and will stay unchanged until the end of the experiment. The mouse enters the box with chamber light off and is left to explore the box with the new element. Whenever the mouse pokes in the nosepoke port, a reward is dispensed. The session ends when the mouse receives 80 rewards or 20 min has elapsed. When 75% of the cohort is retrieving ≥15 pellets during nosepoke shaping phase, the experiment moves to the next phase (usually ~3 days).
- ***SDT– 5 sec Cue Training*.** The mouse enters the box with chamber light off. The session begins with an initial pre-cue delay period of the variable duration of 3-5 sec. If the mouse pokes during this time, the program moves to anticipatory response contingency (see below). Otherwise, it is followed by a 8 kHz pure tone auditory cue (~71 dB) that lasts for up to 5 sec. If the mouse pokes during the cue, the magazine lights up, a chocolate pellet is dispensed and the program moves to ITI contingency (see below). If, on the other hand, the mouse doesn’t poke during the 5 sec cue, the cue turns off and the program moves to post-cue response period that lasts up to 5 sec. If the mouse pokes during this phase, the magazine lights up, a chocolate pellet is dispensed and the program moves to ITI contingency. If, on the other hand, the mouse doesn’t poke during the post-cue response period, the program moves to time out contingency (see below).
- ***SDT– 1 sec Cue Training*.** This phase is exactly as the 5 sec cue training, with the only exception that the tone (cue) stays on for up to 1 sec vs 5 sec. The session ends when either the mouse has reached 100 correct responses or 20 min elapses.

<u>ITI contingency</u>: the magazine light turns off, after a 10 schedule delay, the program returns to pre-cue delay period. If, on the other hand, the mouse pokes during ITI contingency, the program goes to anticipatory response contingency.

<u>Anticipatory response contingency</u>: the chamber light turns on for 10 sec. If the mouse pokes during this time, the program restarts anticipatory response contingency. If, on the other hand, it doesn’t poke, the chamber light turns off and the program moves to pre-cue delay period.

<u>Time out contingency</u>: the chamber light turns on for 10 sec. If the mouse pokes during this time, the program moves to delayed response contingency. If it doesn’t, the chamber light turns off, the trial is considered omitted and the program moves to pre-cue delay period.

<u>Delayed response contingency</u>: the chamber light turns on for 10 sec. If the mouse pokes during this time, the program restarts delayed response contingency. If it doesn’t, the chamber light turns off, the trial is considered omitted and the program moves to pre-cue delay period. Session ends when either the mouse has reached 100 correct responses or 20 min elapses. When 75% of the cohort is getting ≥70% trials rewarded for 2 consecutive days in SDT Training 1, the experiment moves to the next phase.

All SDT experiments were performed within a Habitest Modular Arena and controlled, recorded, and analyzed by Graphic State 4 software (Coulbourn Instruments). were normalized across cohorts and experimental groups relative to 5 second cue training day 1.

#### Attentional Set Shift

One week before the test day, mice started a food deprivation protocol to achieve 80-85% of the initial weight. On day 1 and in each consecutive day, they are handled, weighted and fed ~20gr of food pellet and a few chocolate pellets (Bio-Serv). On the day of the experiment, the mice are introduced in a squared open field arena (16 × 16 × 16 inch) for 5 consecutive trials and their behavior recorded by a camera and analyzed by EthoVision XT (Noldus) software, similar to previous studies^47^. Each of the arena walls has a different visual cue, and in front of each of them, on the floor and ~3 inch from the wall, there is a medicinal cup containing bedding mixed with either sage, or cinnamon, or cumin or cloves (2 gr of spice in 500gr bedding). During the pre-trial (T0), the mice are introduced in the arena for 5 minutes and let explore the cups. This phase was necessary to assess the mice exploratory activity and exclude any odor bias as well as differences between groups in sensitivity to the odors. For each of the successive 4 trials (T1-T4), the mice are re-introduced in the arena for 5 min, and the cup containing sage was enriched by adding a chocolate pellet (reward). From trial to trial, the cups position was randomly shifted so that the odor/visual cue pair was always different, but it was kept fixed for all mice. To correctly perform the task, the mice must learn to ignore the visual cue that remain at a fixed location, and selectively pay attention to the odor as they change position in the maze from trial to trial. During ITI, the mice were moved to a holding cage while the experimenter cleaned the arena with 10% ethanol, replace the cups with clean ones and re-baited the sage cup. The exploration time spent by the mice on each cup was recorded, as well as the latency to reach the correct cup (sage) and retrieve the pellet. Mice that did not locate the chocolate pellet in the initial 3 minutes of trial 1 were excluded from the analysis.

#### Novel Object Recognition Task (NORT)

This test began with 2 days of habituation where the mice are allowed to explore an empty square arena (16’’ × 16’’ × 14’’ (L × W × H) for 15 minutes. During training (day 3), mice are re-introduced in the arena where are now present two identical objects positioned in the back left and right corners of the cage. Each animal was placed in the middle point of the wall opposite to the objects and allowed to explore them for 15 min. At the end of the training phase, mice return to their home cage for 15 min, while the box and the objects were cleaned with 10% Ethanol and then water. Successively, the mice were re-entered into the arena for the test, during which one of the two (familiar) objects was replaced with a new one (novel), totally different in color, texture and shape. Each mouse was left free to explore the objects for 5 min. The entire experiment was recorded using a ceiling mounted camera and the animal behavior was automatically tracked and analyzed by the EthoVision XT (Noldus) software. Two measures were considered: 1) total exploration time (sec) spent by the animal interacting with the two familiar objects during training, in order to evaluate objects bias and 2) the exploration time spent by the animal interacting with the novel object over total exploration time (e.g., [novel/(familiar + novel)] × 100) during the test. Object exploration time is defined as time during which the mouse nose was in contact with the object or directed toward it at a distance ≤ 2 cm.

#### Open Field

Thigmotaxis was determined in an open field box (16 × 16 × 16 inch), virtually divided in a peripheral and a central zone 50% smaller. Each mouse was allowed to explore the apparatus for 15 min and its behavior was recorded by a camera and analyzed by Ethovision. The time spent by the animal in the center of the arena was measured. In this test, the preferential exploration of the peripheral zone of the open field is considered an index of anxiety.

#### Elevated Plus Maze (EPM)

This test is commonly used to evaluate anxiety-like behavior in rodents ^81^. The apparatus was composed of four black plastic arms, arranged as a cross, located 55 cm above the plane of a laboratory bench and illuminated by a 60 w lamp located above the apparatus. Two close arms, opposite to each other were enclosed by lateral walls (50 × 6 × 40 cm), whereas the other two open arms were without walls (50 × 6 × 0.75 cm); the close and open arms delimited a small squared area (6 × 6 cm) called center. Each mouse was placed into the center of the maze, facing one of the two open arms, and its behavior was video-recorded for 5 min and automatically analyzed by the EthoVision XT software (Noldus) for the time spent by the mice in each of the three compartments (open, close, center), which was measured by an observer blind to the experimental groups.

#### 3-Chamber Social Interaction

Tests used a (18’’ × 18’’ × 12’’ (L × W × H) clear acrylic arena, which was divided into 3 chambers of equal area (18” × 6” × 12” (L × W × H) that were separated by walls 6” in length on each side, so that there was a 6” long separation in each wall that a mouse could pass through. Mice were habituated to the testing area for 1 hr prior to the start for the experiment. The test began with a 5 min habituation phase to the center chamber, in which the openings in the walls were obstructed so that the mice could not see or enter either opening. Mice were then put in a transfer cage for 1 minute as the center walls were opened, after which the mice were returned to the center chamber for a 5 min habituation phase to all 3 chambers of the arena. Mice were then returned to the transfer cage for 5 min and the arena as wiped down with 10% ethanol, and wire cups were placed upside down in the center of the outer 2 chambers either with a non-social stimulus (foam figurine) or a novel, age- and strain-matched mouse underneath. Mice were then placed back in the center chamber and allowed to explore for 15 min. Behavior was video-recorded and automatically analyzed by the EthoVision XT Software (Noldus) for time spent in each chamber and time spent exploring a 3 cm proximity to the social or the non-social stimuli (social and non-social zones, respectively). The social discrimination index was calculated as the difference between the mouse’s time in the social zone and the non-social zone, divided by the total time exploring both zones.

#### Go/No-Go behavior

Mice were head-fixed in place above the center styrofoam ball (axially fixed with a rod passing through the center of the ball and resting on post holders) and allowed to move freely forward and backward. MATLAB engine ViRMEn was used to design the virtual task landscape and a National Instruments Data Acquisition (NIDAQ) device provided TTL pulses to trigger the Arduino Unos controlling the tones, odors, airpuff, and lick port. Capacitance changes of the lick port during licking were recorded through the NIDAQ as well.

Prior to behavioral training (2-3 days), homecage water was replaced with water containing 1% citric acid to increase motivation to receive water rewards throughout the task. Habituation began with mice receiving water rewards during Go cues presentation (odor: isoamyl acetate, pure tone: 6kHz). After 3-4 days, mice would be trained using blocks of Go and No-Go cue (odor: lavender oil, pure tone: 1kHz) trials. Delivery of water rewards required mice to lick during Go cues presentation and an aversive airpuff punishment (25 psi) would be delivered the flank of the mouse for licking during No-Go cues. After mice completed the block trials with 70% or greater accuracy, Go and No-Go trials would be pseudo-randomly interleaved (60-80 trials in total).

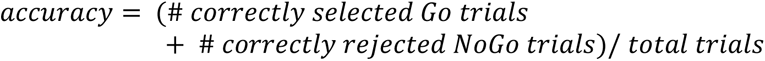

Mice completing the trials with 70% or greater accuracy for two consecutive days would then move on to testing. The testing trial structure is as follows: A 2 second trial start tone (pure tone: 3kHz) begins each trial followed by a 2 second delay then either Go or No-Go cues will be presented (2 second presentation). At the onset of the cue presentation a decision window will begin and last 2.5 seconds. A correct selection of the Go cues is made by licking within this decision window and a water reward is delivered at the end of correctly identified decision periods. Correct rejections of No-Go cues are measured by the absence of licking within the decision window. Each trial is followed by a 15 second inter-trial interval.

#### Head-Fixed Signal Detection Behavior

Following the completion of Go/No-Go testing, mice were tested on a signal detection task. Each trial began with the 2 second trial start tone (pure tone: 3kHz) and following a 2 second delay mice were presented with increasingly shortened Go cues (odor: isoamyl acetate, pure tone: 6kHz; cue length: 2 seconds, 1 second, 0.5 seconds). After the Go cues presentation began, a decision window of 2.5 seconds opened and mice that licked within this window received a water reward.

### QTL mapping in Diversity Outbred Mice

#### Genotype Identification & Haplotype Reconstruction

SNP locations and genotypes for the eight founder strains were acquired from ftp.jax.org/MUGA and the consensus genotype for each founder strain and each SNP was determined from the multiple individuals that were genotyped. SNP genotypes for the 182 DO mice were determined using a high density mouse universal genotyping array, GigaMUGA (geneSeek). A total of 114,184 SNPs were detected on the 19 autosomes and X chromosomes. Using R/qtl2 ^35^, founder haplotype probabilities were reconstructed for all samples and then converted to additive allelic dosages and scaled to 1. Realized genetic relationship matrices, often referred to as kinship matrices, were estimated using the leave one chromosome out (LOCO) method, so that the kinship term does not absorb variation explained by the putative QTL. Another QTL mapping software package for multi-parental populations (MPP), miQTL, was used to confirm findings from R/qtl2, and to visualize and assess the level of heterozygosity at the locus of interest.

#### QTL Mapping

Phenotype values from the prepulse inhibition performance were subject to Box-Cox transformation. Then, using R/qtl2, an additive single locus linear mixed model was fit at positions across the genome, producing a genome scan. Potential population structure was controlled for through the inclusion of a random effect to account for correlation structure measured by the kinship matrix. This was performed in R/qtl2 using the leave one chromosome out (LOCO) method ^82^. For confirmation of the QTL results, we performed a multiple imputation genome scan (11 imputations) using miQTL, to assess whether uncertainty in founder haplotype reconstruction was strongly influencing the results. Genome-wide significance thresholds (alpha = 0.05) for the genome scans were determined through 1000 permutations of the diplotype.

#### Analysis of Founder Contributions

To determine the founder haplotype effects driving the Chr13QTL, we first estimated best linear unbiased predictors (BLUPs), which constrain potentially unstable effects by fitting the QTL term as a random effect. To further confirm these results, we used Diploffect, to estimate posterior credible intervals for the haplotype effects as well as the proportion of variance explained by the QTL (sometimes referred to as the locus heritability).

### RNA Expression Analysis

#### RNA Extraction From Brain Tissues

For tissue extraction, p28 and adult (up to p120) mice were sacrificed by cervical dislocation and immediately decapitated, while p7, p14, and p21 mice were sacrificed by decapitation in compliance with IACUC protocol # 22087-H. The targeted brain regions were harvested from 1 mm brain slices, obtained by brain matrices (ZIVIC) using 1.0 mm tissue punches and transferred to a tube containing 300 mL of ice-cold lysis buffer and 3 mL ß-mercaptoethanol (Total RNA Purification kit, NORGEN; following the manufacturer’s protocol). Samples were then homogenized by passing a 25G insulin syringe six times and left on ice. For RNA extraction, the total RNA Purification kit was used according to the manufacturer’s instructions (NORGEN). RNA quality was evaluated by Bioanalyzer 2100 (Eukaryote Total RNA Nano chip, Agilent) at the Rockefeller University Genomic Resource Center (RIN > = 7.50 and free of genomic DNA contamination). RNA samples were then aliquoted and stored at −80°C.

#### Bulk RNA Sequencing (RNA-seq) and Analysis

For RNAseq, RNA libraries were prepared from 100ng of total RNA per sample for 6 DO mice, 3 brain regions per mouse using the TruSeq stranded mRNA LT kit (Cat# RS-122-2101, Illumina). These synthetic RNAs cover a range of concentrations, length, and GC content for validation of the fidelity and dose-response of the library prior to downstream procedures. Libraries prepared with unique barcodes were pooled at equal molar ratios following manufacturer’s protocol (Cat# 15035786 v02, Illumina). The pool was denatured and subject to paired-end 50x (DO samples) or single-end 100x (CC samples) sequencing on the NovaSeq SP platform. An average of 67 million reads per sample were obtained. Sequencing reads were aligned to the mouse genome (mm10) using STAR (v2.4.2a) and aligned reads were quantified using Salmon (v0.8.2). Approximately 90% of the reads mapped uniquely. Hierarchical clustering and Principal Components Analysis were performed following Variance Stabilizing Transformation (VST) from DESeq2, which is on the log2 scale and accounts for library size differences. The hierarchical clustering heatmap shows the Euclidean distances of VST of the counts between samples.

#### Quantitative PCR (qPCR)

For quantitative PCR, each reverse transcription was performed with 0.2 mg RNA using the High-Capacity RNA-to-cDNA kit (Applied Biosystems # 4387406), in a final volume of 20 µL. Primers for reverse transcription were equal mixtures of poly-T nucleotides and random hexamers. Negative controls (omitting reverse transcriptase enzyme) were performed for each sample. The cDNA products were diluted 1:1 and 2 µL was analyzed by qPCR using custom primer sets and PowerUp SYBR Green Master Mix (10 µL total reaction, Applied Biosystems # A25742). RT-qPCRs were performed on a Quantstudio3 from Applied Biosystems. Every reaction was systematically run in triplicate. Conditions were the following: 50°C 2 min, 95°C 10 min, 40 × (95°C 15 s, 60°C 1 min). qPCR Ct values were analyzed using the LightCycler software. Detection threshold was set at DRn = 0.3, with this limit always within the 2n exponential amplification phase of genes. Mean of technical triplicate values were reported. All mice gene expression Ct values were normalized with the reference gene Ube2d2a using dCt method to determine the relative mRNA expression of each gene. Developmental knockdown mice that expressed both *Homer1a* and *Ania3* at levels higher than the average scramble expression by half a standard deviation or more were post hoc excluded from downstream analyses.

### Single-Cell Sequencing

#### Single-Cell Dissociation and Single-Cell RNA Sequencing

Single cell suspensions of prefrontal cortex were prepared as described previously ^48^. Briefly, mice were sacrificed with an overdose of isoflurane, followed by transcardial perfusion with carbogenated (95% O_2_, 5% CO_2_) Hanks’ Balanced Salt Solution (HBSS). Brains were removed, 500μm sections were collected, and the prefrontal cortex region was isolated. The tissue was dissociated using papain (LS003124, Worthington) dissolved in Hibernate A buffer (NC1787837, Fisher Scientific) and incubated for 25-30 min at 37°C, followed by manual trituration using fire polished Pasteur pipettes and filtering through a 40μm cell strainer (BAH136800040, Millipore Sigma). Cells were washed with wash buffer (PBS + 1% BSA) and centrifuged at 200 g for 5 min, the supernatant was carefully removed, and cells were resuspended in ~500ul wash buffer and 10% DAPI. Flow cytometry was done using a BD FACS Aria III Cell Sorter (BD FACSDiva Software, v8.0.1) with a 100-µm nozzle. The cell suspensions were first gated on forward scatter, then within this population based on lack of DAPI staining. Cells were collected in wash buffer, manually counted using a Burker chamber, and suspension volumes were adjusted to a target concentration of 700 − 1000 cells/μl. Single cell RNA-sequencing was carried out with the Chromium Next GEM Single Cell 3’ Kit v3.1 (10X Genomics, 1000268). Manufacturer’s instructions were followed for downstream cDNA synthesis (12-14 PCR cycles) and library preparation. Final libraries were sequenced on the Illumina NovaSeq S4 platform (R1 – 28 cycles, i7 – 8 cycles, R2 – 90 cycles).

#### Single-Cell RNA Sequencing Data Analysis

RAW sequencing reads were aligned to the GRCm38/mm10 mouse reference genome and a custom cell-gene count matrix was constructed using the Sequence Quality Control (SEQC) package ^83^. Viable cells were identified on the basis of library size and complexity, whereas cells with >20% of transcripts derived from mitochondria were excluded from further analysis. The Python Scanpy package (v1.9.3) was used to further analyze the data. Replicates were merged and doublets were removed using Scrublet ^84^. Cells with <2,500 UMIs per cell and <1,000 genes per cell, and genes detected in <3 cells were removed. Per-cell counts were normalized to equal the median of total counts per cell, and log-transformed. Principal component analysis was used to reduce the dimensionality to 50 principal components. A nearest neighbor graph was computed between cells using these principal components, and Leiden clustering was applied to separate the cells into distinct clusters of major cell types. Known gene markers were used ^48^ to assign cell types. Once the neuronal cluster was identified, it was subsetted and re-clustered using the first 50 principal components to identify inhibitory and excitatory neurons. Clusters with differential *Homer1* expression between cc083 and cc025 strains were identified using t-tests. Clusters with significantly different *Homer1* expression between strains were merged, and the “MAST” R package ^52^ was used to identify differentially expressed genes between strains for the merged cluster as well as all individual clusters. Gene set enrichment analysis was performed using the fast gene set enrichment analysis (GSEA) package (fgsea v1.18.0), the GO_Molecular_Function_2021 gene set, and the <u>Elsevier_Pathway_Collection</u> gene-set libraries using Enrichr ^53–55^.

### Gene Expression Maipulation Experiments In Vitro & In Vivo

We used the following shRNAs for gene knockdown (which were then subcloned into a pscAV-U6-mCherry construct, VectorBuilder/Vector Biolabs):

Homer1a (GenBank: NM_011982.4), Targeting sequence: GGTTTCAGAAACTCTTGAA; Ania3 (GenBank: NM_001347598.1),Targeting sequences: GGAGACATAGTTCTTCTTA, GCTAAGCTAGAGCCATCTA.

For gene expression, coding sequences of Homer1a and Ania3 was cloned from mouse cortical cDNA and subsequently subcloned into a pAAV.CamKII(1.3).eYFP.WPRE.hGH expression vector using standard molecular cloning techniques. Constructs were verified first by Sanger sequencing, and then diagnostics for ITR integrity, by digestion with SmaI, before AAV production.

### Generation of AAV-MAG-GCaMP6f

We identified the mouse *MAG* gene locus (including introns and a 3 kb upstream potential promoter region) using the UCSC genome browser, as others have done previously ^85^, on the reverse strand of Chr 7qB1: 30,899,176-30,917,832 in the July 2007 mm9 alignment (Chromosome 7: 30,598,601-30,617,298 in the GRC38/mm10 alignment). Sequence conservation was assessed using the VISTA genome browser ^86^ and the putative MAG promoter was screened for regions of >50% interspecies sequence similarity, which were then evaluated for transcription factor binding sites, especially OL-lineage specific Olig1 and Olig2, using the Wilmer Bioinformatics Resource ^87^ and the P-match 1.0 program (http://gene-regulation.com/pub/programs.html#pmatch). This method yielded a 2.5 Kb putative MAG promoter region. The putative MAG promoter was cloned from mouse cortical cDNA using standard molecular cloning techniques and replaced the Syn promoter from the pAAV.Syn.GCaMP6f.WPRE.SV40 plasmid. The pAAV-MAG-GCaMP6f construct was packaged using the Olig001 capsid ^79^, which has high oligodendrocyte tropism. (Univ. of Arizona Viral Production Core).

### Histology & Immunohistochemistry

Mice were transcardially perfused with PBS and 4% paraformaldehyde in 0.1M PB, then brains were post-fixed by immersion for 24 h in the perfusate solution followed by 30% sucrose in 0.1M PB at 4°C. The fixed tissue was cut into 40 mm coronal sections using a freezing microtome (Leica SM2010R), stained with DAPI (1:1000 in PBST), and mounted on slides with ProLong Diamond Antifade Mountant (Invitrogen). For immunostaining, the fixed sections were permeabilized with 70% methanol for 15 min before blocking with 5%normal donkey serum in PBS for 1 h and incubated with primary antibodies overnight at 4°C. Sections were washed three times in PBS and incubated with appropriate secondary antibodies overnight at 4°C. Afterward, coverslips were mounted using ProLong Diamond Antifade mounting medium for image collection. Primary and secondary antibodies include rabbit polyclonal anti-NeuN (Millipore ABN78), rabbit polyclonal anti-Iba1 (Wako, 019-19741), rabbit polyclonal anti-Olig2 (Millipore, AB9610), and mouse monoclonal anti-GFAP (Millipore MAB360), Alexa Fluor 647 donkey anti-rabbit IgG (Jackson ImmunoResearch, Cat # 711-606-152), Alexa Fluor 647 donkey anti-mouse IgG (Jackson ImmunoResearch, Cat # 715-606-151), and DAPI (Cayman Chemical, Cat#28718-90-3). For immunohistochemistry staining, epifluorescent images were obtained at room temperature on a Nikon Eclipse T*i* microscope using a Nikon 4x (NA 0.13, dry), 10x (NA 0.30, dry), or 20x (NA 0.45, dry), objectives with the same settings and configurations for each objective across all samples within each experiment.

### In Vivo Multi Site Photometry Recordings

#### Photometry Setup

A custom dual-color, multi-fiber photometry setup was built. For GCaMP6f imaging, excitation of the 470 nm (imaging) and 405 nm (isosbestic control) wavelengths were provided by LEDs (Thorlabs M470F3, M405FP1), which were collimated into a dichroic mirror holder with a 425 nm long pass filter (Thorlabs DMLP425R). This was coupled to another dichroic mirror holder with a 495 nm long pass dichroic (Semrock FF495-Di02-25x36) which redirected the excitation light on to a custom branching low-autofluorescence fiberoptic patchcord of three bundled 400 mm diameter 0.57NA fibers (BFP(3)_400/440/PKMJ-0.57_1m_SMA-3xFC_LAF, Doric Lenses) using a 20x/0.5NA Objective lens (Nikon CFI SFluor 20X, Product No. MRF00100). GCaMP6f fluorescence from neurons below the fiber tip in the brain was transmitted via this same cable back to the mini-cube, where it was passed through a GFP emission filter (Semrock FF01-520/35-25), amplified, and focused onto a high sensitivity sCMOS camera (Prime 95b, Photometrics). For jRGECO1a imaging, a second light path was built so that excitation of the 565 nm (imaging) and 470 nm (isosbestic control) wavelengths were provided by LEDs (Thorlabs M565F3^h^, M470F3), which were collimated into a dichroic mirror holder with a 505 nm long pass dichroic (Thorlabs DMLP505R). This was coupled to another dichroic mirror holder with a 573 nm long pass dichroic (Semrock Di02-R561-25x36), which redirected the excitation light on to a low-autofluorescence monofiberoptic patchcord with a 400 mm diameter 0.57NA fiber (MFP_400/440/PKMJ-0.57_1m_SMA-FC_LAF, Doric Lenses) using a 20x/0.5NA Objective lens (Nikon CFI SFluor 20X, Product No. MRF00100). jRGECO1a fluorescence from neurons below the fiber tip in the brain was transmitted via this same cable back to the mini-cube, where it was passed through a RFP emission filter (Semrock FF01-607/36-25), amplified, and focused onto a high sensitivity CMOS camera (BFS-PGE-50S5M-C, Teledyne FLIR).

Each of the multiple branch ends of the branching fiberoptic patchcord as well as the monofiberoptic patchchord were coupled to four 2 m low-autofluorescence patchcords (MFP_400/430/1100-0.57_2m_FCZF1.25_LAF, Doric Lenses) which is used to collect emission fluorescence from 1.25mm diameter light weight ferrules (MFC_400/430-0.48_ZF1.25, Doric Lenses) using a mating sleeve (SLEEVE_BR_1.25, Doric Lenses). An microcontroller (Arduino Uno) was programmed to take trigger inputs from the Operant Behavior Setup or MATLAB and synchronize the camera shutters and alternate triggering of the 405 nm and 565 nm LEDs together and both 470 nm LEDs together. Custom TTL converters were used to read in frame acquisition times to the Habitest Modular system (above), which were integrated with events from the behavior in Graphic State 4. Bulk activity signals were collected using the PVCAM (GCaMP) and Spinnaker (jRGECO) software, and data were further post-processed and analyzed using custom MATLAB scripts.

### Quantification and Statistical Analysis

#### Behavior Statistical Reporting

Sample sizes were selected based on expected variance and effect sizes from the existing literature, and no statistical methods were used to determine sample size a priori. Prior to experiments being performed, mice were randomly assigned to experimental or control groups. The investigator was blinded to all behavioral studies (except for CC083 versus CC025 cohorts, where coat color differences prevent blinding during experimentation). Homer1a/Ania3 shRNA mice were removed from the developmental knockdown experiments if they did not have sufficiently reduced expression relative to the scramble groups or were extreme outliers from the remainder of the knockdown mice. Data analyses for calcium imaging were automated using MATLAB scripts. Statistical tests were performed in MATLAB 2022b or Graphpad Prism 9.

#### Gene Expression Statistics

Differential gene expression between high and low performing DO mice as well as between CC025 and CC083 mice from bulk RNAseq data was determined in R (3.5.0) using the DESeq2. P values were determined using a Wald test and p values were corrected using the Benjamini-Hochberg (BH) method.

#### Multi-Fiber Photometry Data Processing

For analysis, the images captured by the sCMOS camera were post-processed using custom MATLAB scripts. Regions of interest were manually drawn for each fiber to extract fluorescence values throughout the experiment. The 405 nm (GCaMP) or 470 nm (jRGECO) reference traces were scaled to best fit the 470 nm (GCaMP) or 565 nm. (jRGECO) signal using least-squares regression. The normalized change in fluorescence (dF/F) was calculated by subtracting the scaled 405 nm or 470. nm reference traces from the 470 nm or 565 nm signals, respectively, and dividing those values by the scaled reference traces. The true baseline of each dF/F trace was determined and corrected by using the MATLAB function ‘‘msbackadj’’ estimating the baseline over a 200 frame sliding window, regressing varying baseline values to the window’s data points using a spline approximation, then adjusting the baseline in the peak range of the dF/F signal. Task events (e.g., cue on/offsets, and nosepokes), were time stamped via the Graphic State 4 software.

#### Multi-Fiber Photometry Data Analysis

Total mean activity, for different task phases, and different strains, were quantified as area under the curve (AUC) of dF/F responses shifted above 0. AUC was calculated using MATLAB “trapz” function and normalized with the recorded time. Pearson Correlation of the dF/F responses was performed between different regions using the ‘‘corr’’ (MATLAB) function. To ensure that correlation values were significantly more than chance, each timeseries was scrambled 10,000 times randomly, for each session across all mice. All such chance correlation coefficients were pooled to calculate mean (all of which were at or near zero) and standard deviation of chance correlations. To quantify signal to noise ratio (SNR), we calculated the mean and standard deviation of each region’s neural activity (z-scored dF/F) during baseline periods of the task, ie all omission trials (from cue onset to the onset of the pre-trial delay phase, calculated values referred to as mean-baseline and SD-baseline) for each mouse for a given day. Trial SNR was calculated as the difference between the maximum pre-cue activity (z-scored dF/F for the 5 seconds immediately prior to cue onset) and the mean-baseline value for that mouse, divided by the SD-baseline value (i.e. (trial pre-cue maximum − mean-baseline) / SD-baseline). For cohorts that progressed to the 1 second cue training phase, only mice remaining above 70% performance accuracy were include in the analyses. Additionally, the first training session and any training sessions under 15 minutes long were not included in the analyses.

---

**Figure S1.**
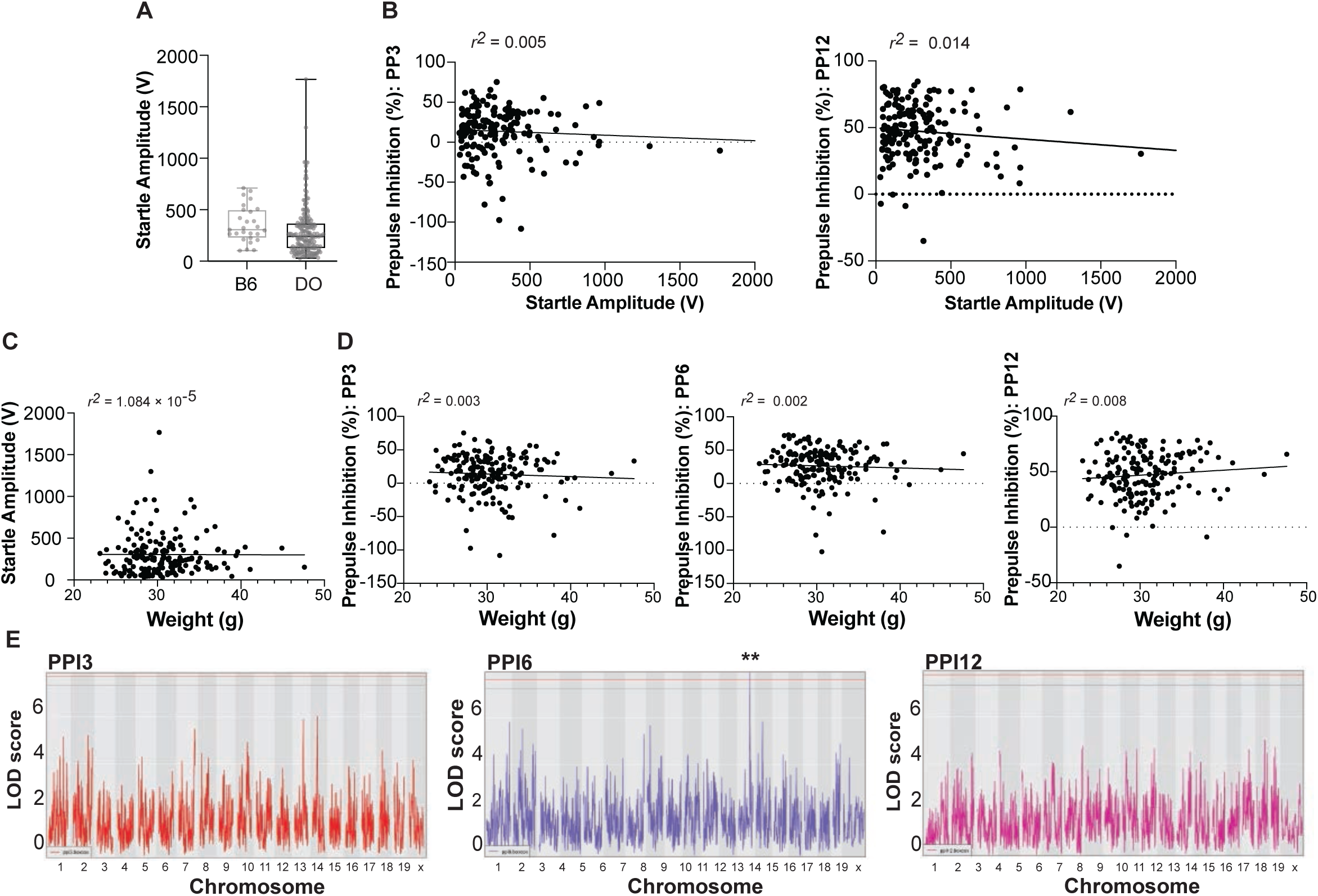
Additional genetic and behavioral characterization of DO mice, related to Figure 1. (**A**) Startle response assessed during PPI experiments in B6 (grey, n=27) and DO (black, n=176) mice measured as startle amplitude (V). Boxes indicate 2nd and 3rd quartiles with median and range. (**B-D**) Correlations in DO mice (n=176) between (**B**) startle response, measured as the magnitude of startle amplitude (V), and PPI, measured as percent inhibition, at 3 (PP3, r^2^=0.005) and 12 (PP12, r^2^=0.014) dB above background, (**C**) weight and startle response (r^2^=1.084 × 10-5), and (**D**) weight and PPI (PP3, r^2^=0.003; PP6, r^2^=0.002; PP12, r^2^=0.008) dB above background. (**E**) QTL mapping analysi s (by R/qtl2), shown as Manhattan plots, of PPI at 3 (PPI3, red), 6 (PPI6, purple, genome-wide p<0.01), and 12 dB (PPI12, magenta) above background (n=176; blue lines indicate 90% confidence threshold and red lines indicate 95% confidence threshold).

**Figure S2.**
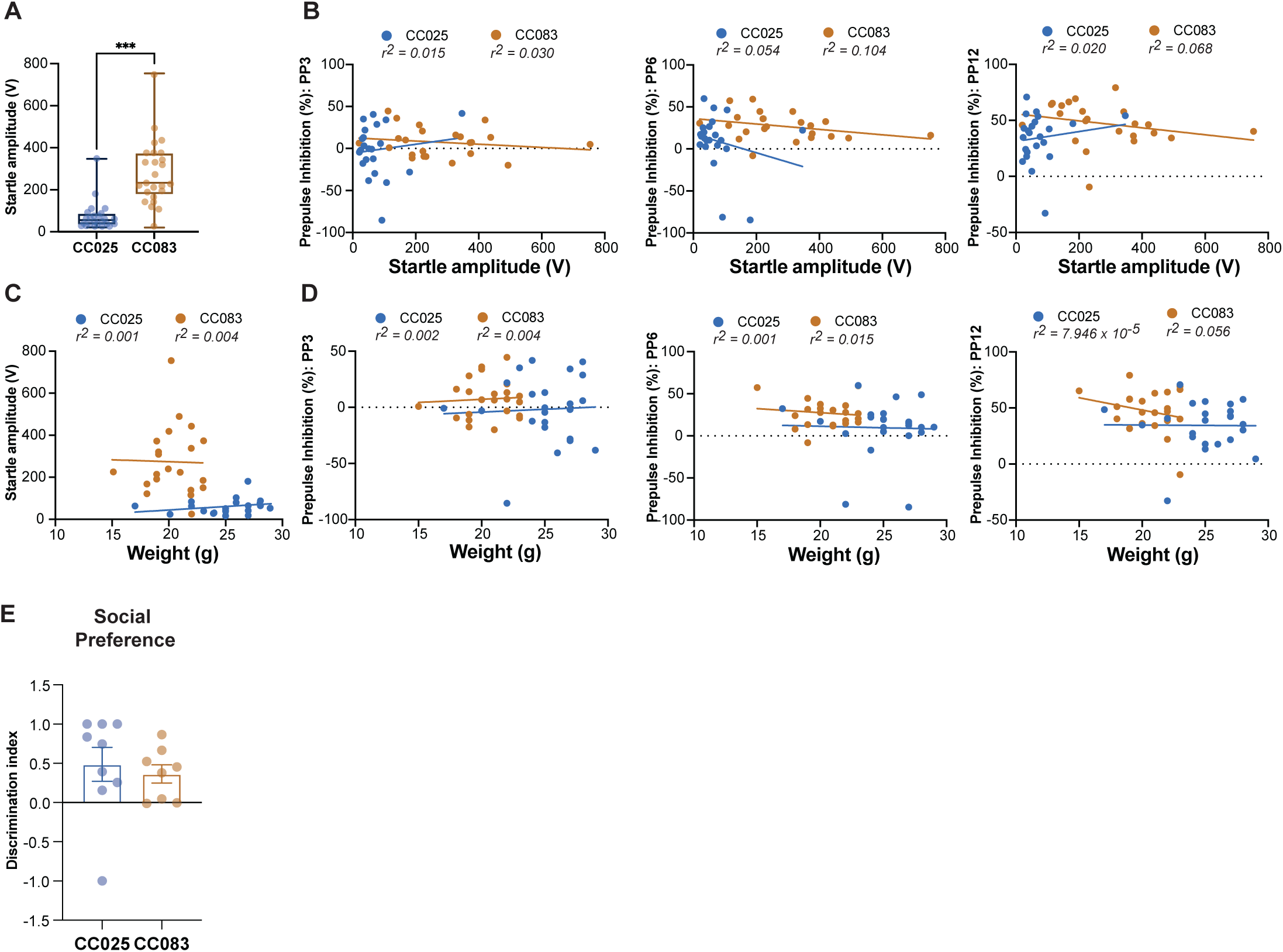
Behavioral phenotype and covariate characterization of CC083 and CC025 mice, related to Figure 2. (**A**) Startle response, measured as the magnitude of the startle amplitude (V) in CC025 (blue, n=24) and CC083 (tan, n=27) mice. Boxes indicate 2nd and 3rd quartiles with median and range. (**B-D**) Correlations in CC025 (n=24) and CC083 (n=27) between (**B**) startle response and PPI, measured as percent inhibition, at 3 dB (PP3: CC025 r2=0.015, CC083 r2=0.030), 6 dB (PP6: CC025 r2=0.054, CC083 r2=0.104), and 12 dB (PP12: CC025 r2=0.020, CC083 r2=0.068) above background, (**C**) weight and startle response (CC025 r2=0.001, CC083 r2=0.004), and (**D**) weight and PPI (PP3: CC025 r2=0.002, CC083 r2=0.004; PP6: CC025 r2=0.001, CC083 r2=0.015; PP12: CC025 r2=7.946 × 10-5, CC083 r2=0.056). (**E**) Social behavior for CC025 (n=9) and CC083 (n=8) mice, expressed as discrimination index determined by exploration time in a 3-chamber social interaction test, shown as mean ± SEM.

**Figure S3.**
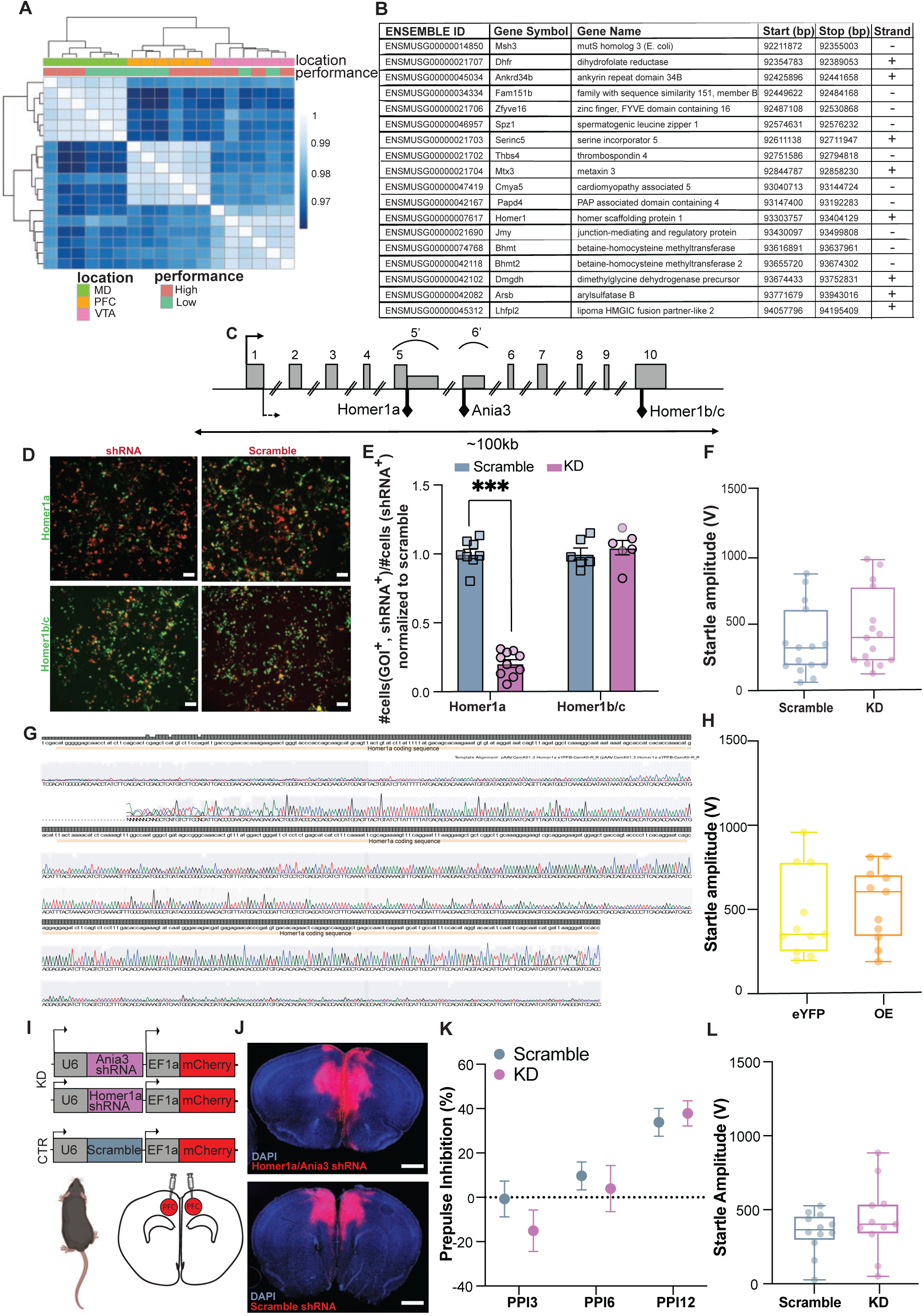
DO RNAseq clustering information, Homer1 exon structure, *in vitro* validation of *Homer1a* knockdown and over-expression vectors, & additional behavioral characterization of adult knockdown & overexpression manipulations, related to Figure 3. (**A**) Heatmap of hierarchical clustering by Euclidean distance among gene expression profiles in DO high-(pink, n=3) and low-performers (green, n=3) as highlighted in Figure 3A-B and from three brain regions per mouse: mediodorsal thalamus (MD, green), prefrontal cortex (PFC, orange) and ventral tegmental area (VTA, pink). Clustering is visible by brain region and performance in MD and PFC. (**B**) Table showing protein-coding genes within the 95% CI surrounding the Chr 13 QTL identified by rQTL2. (**C**) Schematic representation of the *Homer1* genomic exon structure. The bent arrow at the 5’ end of exon 1 (solid line, above) indicates the putative transcription start site, while the bent arrow at the 3’ end of exon 1 (dashed line, below) represents the translation start site. Black diamonds (below) indicate the translation stop sites of *Homer1a*, *Ania3*, and *Homer1b/c,* respectively. To create *Homer1a*, exon 5 extends into intron 5 to create the *Homer1a*-specific exon (5’) through alternative splicing. *Ania3* is generated by alternative splice usage of intron 5 sequence downstream of exon 5’ as the *Ania3*-specific exon 6’. (Adapted from Bottai et al. 2002). (**D-E**) *in vitro* validation of *Homer1a* gene knockdown construct. (**D**) Representative images of HEK cells co-transfected with *Homer1a* (left column) or Scramble (right column) shRNA (red) and *Homer1a* (top row) or *Homer1b/c* (bottom row) expression constructs (green). Scale bar: 100 µm. (**E**) Quantification of shRNA-mediated gene knockdown, expressed as the fraction of cells co-expressing a *Homer1* isoform construct and shRNA construct relative to the total number cells expressing the shRNA construct, normalized to the respective scramble control experiments (two-way ANOVA showed significant main effects for *Homer1* isoform expression, p<0.001, and shRNA construct, p<0.001, as well as a significant interaction between those variables, p<0.001; Holm-Sidak’s test for multiple comparisons showed a significant difference in *Homer1a* expression between the shRNA (purple, n=10 fields of view across 2 independent experiments) and Scramble (blue, n=8 fields of view across 2 independent experiments) constructs, p<0.001). (**F**) Startle response in Scramble (n=15) and adult KD (n=15) mice. Boxes indicate 2nd and 3rd quartiles with median and range. (**G**) Electropherogram of AAV-CaMKII(1.3)-Homer1a-eYFP overexpression construct aligned to the *Homer1a* coding sequence (tan bar near the top of each line). The height of grey boxes at the top of each line is proportional to the number of sequencing runs aligned to the reference sequence (maximum # of sequencing runs in image = 2). (**H**) Startle response in control eYFP (yellow, n=15) and OE (orange, n=15) mice. Boxes indicate 2nd and 3rd quartiles with median and range. (**I**) Schematic of constructs and injection location (PFC) for knockdown (KD, purple) and control (Scramble, blue) in adult B6 mice. (**J**) Validation histology performed 12 weeks after bilateral injection of pooled AAV-U6-Homer1a shRNA-EF1A-mCherry and AAV-U6-Homer1a shRNA-CMV-mCherry and AAV-U6-Ania3 shRNA-EF1A-mCherry for KD (purple, upper panel) and AAV-U6-Scramble-EF1A-mCherry control virus for Scramble (blue, lower panel) into PFC, showing viral transduction in the target area (DAPI, blue; mCherry, red). Scale bars: 1000 µm. (**K**) PPI in Scramble (n=12) and adult *Homer1a/Ania3* KD, (n=11). (**L**) Startle response in Scramble (n=12) and adult KD (n=11) mice. Boxes indicate 2nd and 3rd quartiles with median and range. Data in **E**, **H**, and **K** expressed as mean ± SEM.

**Figure S4.**
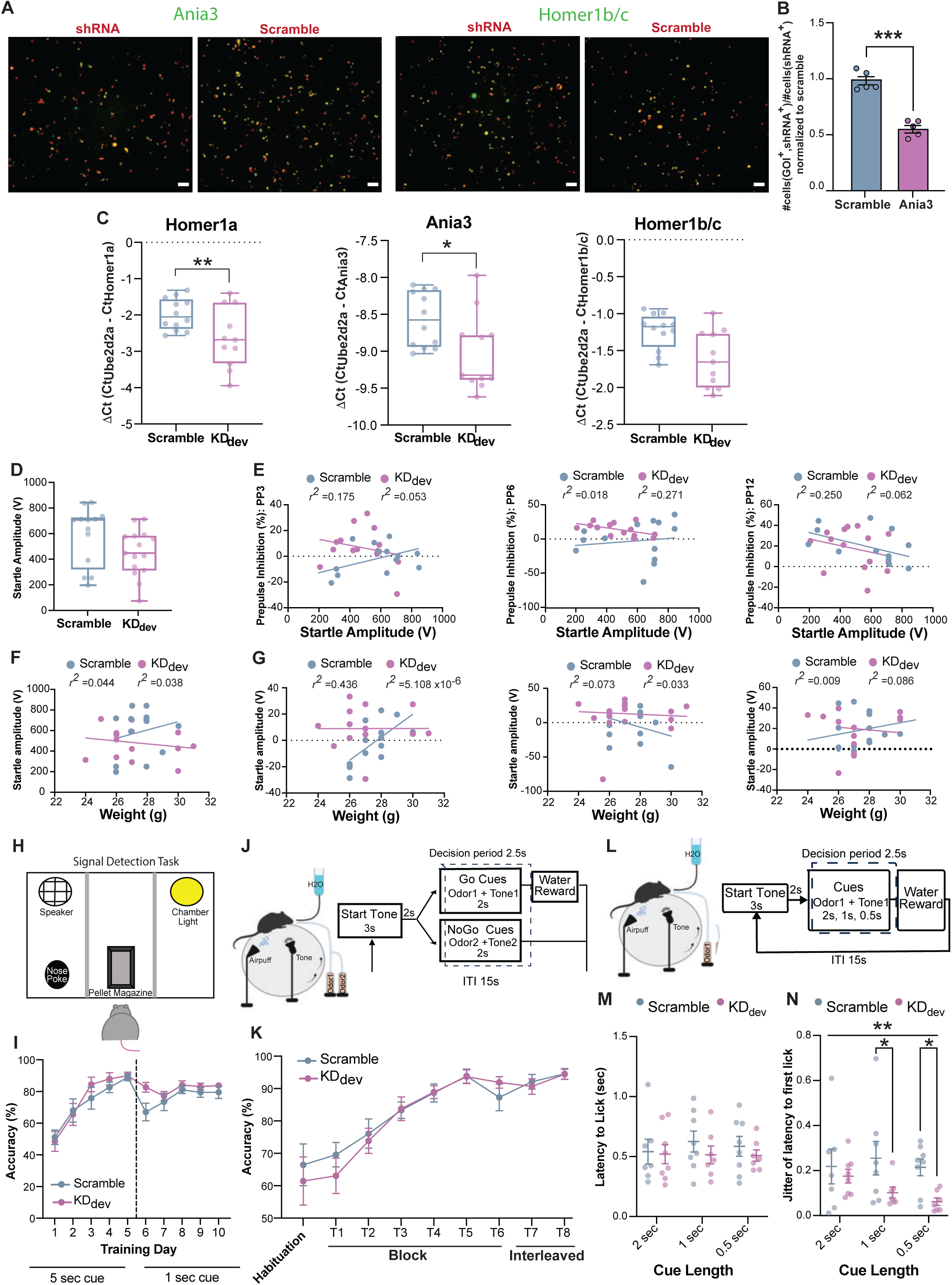
*In vitro* characterization of Ania3 shRNA construct and *in vivo* characterization of developmental knockdown (KD_dev_) manipulation, related to Figure 4. (**A**) *in vitro* validation of gene knockdown constructs. Representative images of HEK cells co-transfected with *Ania3* (panels 1 and 3 from the left) or Scramble (panels 2 and 4 from the left) shRNA (red) and *Ania3* (panels 1 and 2 from the left) or *Homer1b/c* (panels 3 and 4 from the left) expression constructs (green), Scale bar: 100 µm. (**B**) Quantification of shRNA-mediated gene knockdown, expressed as the fraction of cells co-expressing the *Ania3* expression construct and shRNA or scramble construct relative to the total number cells expressing the shRNA or scramble, normalized to the scramble control experiments. In cells transfected with the *Ania3* expression construct, there was a significant difference in *Ania3* expression between the cells co-transfected with the shRNA (n=5 fields of view), and Scramble (n=5 fields of view) constructs (Unpaired t-test, p<0.001). (**C**) *ex vivo* validation of developmental knockdown manipulation assessed by quantification of *Homer1a* (left), *Ania3* (center) and *Homer1b/c* (right) levels measured by qPCR in PFC samples dissected from Scramble (n=12) and KD_dev_ (n=15), (two-way ANOVA showed significant main effects for group, p<0.001, and *Homer1* isoform expression, p<0.001; post hoc Holm-Sidak’s test for multiple comparisons shows a significant difference in *Homer1a,* p=0.004, and *Ania3,* p=0.05, expression). (**D**) Startle response in KD_dev_ (n=15) and Scramble (n=13) mice, measured as the magnitude of startle amplitude. (**E-G**) Correlations between (**E**) startle response and PPI, measured as percent inhibition, at 3 dB (PP3: Scramble r^2^ = 0.175, KD_dev_ r^2^ = 0.053,), 6 dB (PP6: Scramble r^2^ = 0.018, KD_dev_ r^2^ = 0.271,), and 12 dB (PP12: Scramble r^2^ = 0.250, KD_dev_ r^2^ = 0.062,) above background, (**F**) weight (g) and startle response (Scramble r^2^ = 0.044, KD_dev_ r^2^ = 0.038,), and (**G**) weight and PPI (PP3: Scramble r^2^ = 0.436, KD_dev_ r^2^ = 5.108 x10-6,; PP6: Scramble r^2^ = 0.073, KD_dev_ r^2^ = 0.033; PP12: Scramble r^2^ = 0.009, KD_dev_ r^2^ = 0.086). (**H**) Schematic of operant wall of arena used for signal detection task (SDT). (**I**) Percentage of correct responses for Scramble and KD_dev_ mice (5 sec cue: n=13 per group; 1 sec cue: Scramble n=9 KD_dev_ n =11). (**J**) Schematic of Go/No-Go task setup (left) and the task structure for testing day 1 (right). (**K**) Go/No-Go task performance accuracy across habituation and training days (n = 8 per group). (**L**) Diagram of head-fixed SDT setup (left) and task structure (right). (**M**) Quantification of the latency to first lick (sec) within the decision windows across cue lengths. Each point is the average latency to first lick for the first 3 Go trials per animal (2 sec cue: Scramble n=7, KD_dev_ n=8; 1 sec and 0.5 sec cues: Scramble n=8, KD_dev_ n=7). (**N**) Quantification of the latency to first lick jitter across cue lengths. Jitter is quantified as the standard deviation of first lick latencies across the first 3 Go trials (two-way ANOVA showed a significant main effect for group, p=0.007, and post hoc Holm-Sidak’s test for multiple comparisons showed significant differences between groups at 1 and 0.5 sec cues, p=0.04 for both cue lengths, 2 sec cue: Scramble n=7, KD_dev_ n=8; 1 sec and 0.5 sec cues: Scramble n=8, KD_dev_ n=7). Data in **C**, **I**, **K**, **M**, and **N** are expressed as mean ± SEM.

**Figure S5.**
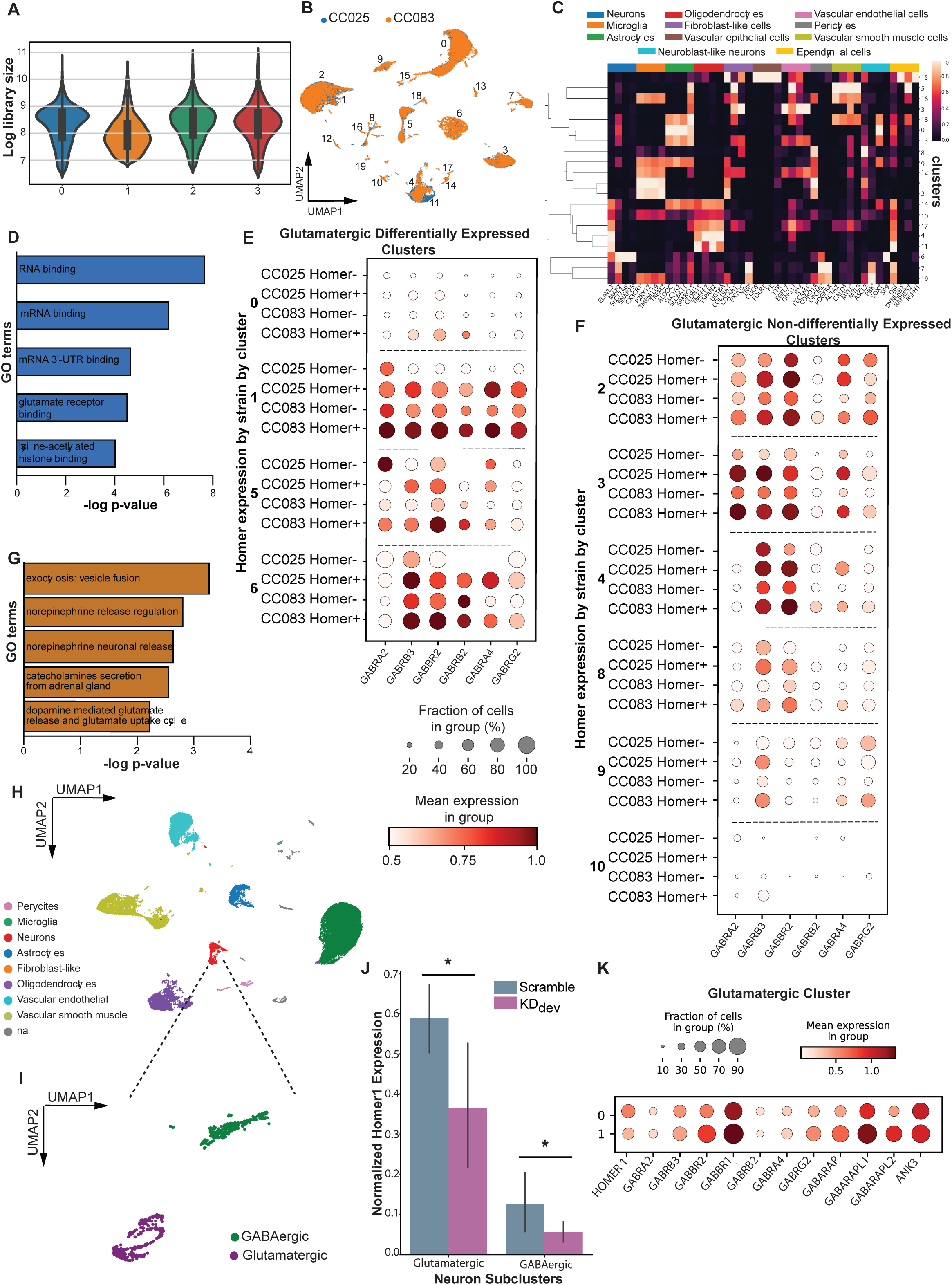
Additional information for scRNAseq experiments, related to Figure 5. (**A**) Violin Plots of library size for each biological replicate (0=pooled CC025 sample 1, 1=pooled CC083 sample 1, 2=pooled CC025 sample 2, 3=pooled CC083 sample 2). (**B**) UMAP visualization of initial clusters colored by line. (**C**) Heatmap of select cell type marker genes for clusters shown in **B**. (**D**) Gene ontology (GO) analysis of molecular function for genes upregulated in CC025 cells within the glutamatergic *Homer1* differentially expressed (DE) clusters. (**E-F**) Dot plots showing scaled expression of GABAergic receptors driving GO analysis (Fig. 5g) in both the glutamatergic *Homer1* (**E**) DE and (**F**) non-differentially expressed (NDE) clusters stratified by cluster, line, and *Homer1* positivity. The size of each dot corresponds to the percentage of cells expressing each gene and color intensity indicates relative, scaled expression of that gene. (**G**) Functional pathway enrichment analysis for CC083 cells in the GABAergic cluster using the Elsiver_Pathway_Collection gene set library in Enrichr. (**H**) UMAP visualization of all cells collected from Scramble and KD_dev_ mice (n=3 mice pooled per group) clustered based on transcriptional profile. (**I**) UMAP visualization sub-clustering all cells identified as neurons, identified as excitatory (glutamatergic) and inhibitory (GABAergic) neuron clusters based on expression of canonical marker genes. (**J**) Differential *Homer1* expression between Scramble and KD_dev_ neurons by cluster (p=0.03 for both glutamatergic and GABAergic clusters). Data shown as mean ± SD. (**K**) Dot plots showing scaled expression of select GABAergic receptors in the glutamatergic cluster by group. The size of each dot corresponds to the percentage of cells from each group expressing each gene or gene set, and the color intensity indicates the relative, scaled expression of the gene/gene set.

**Figure S6.**
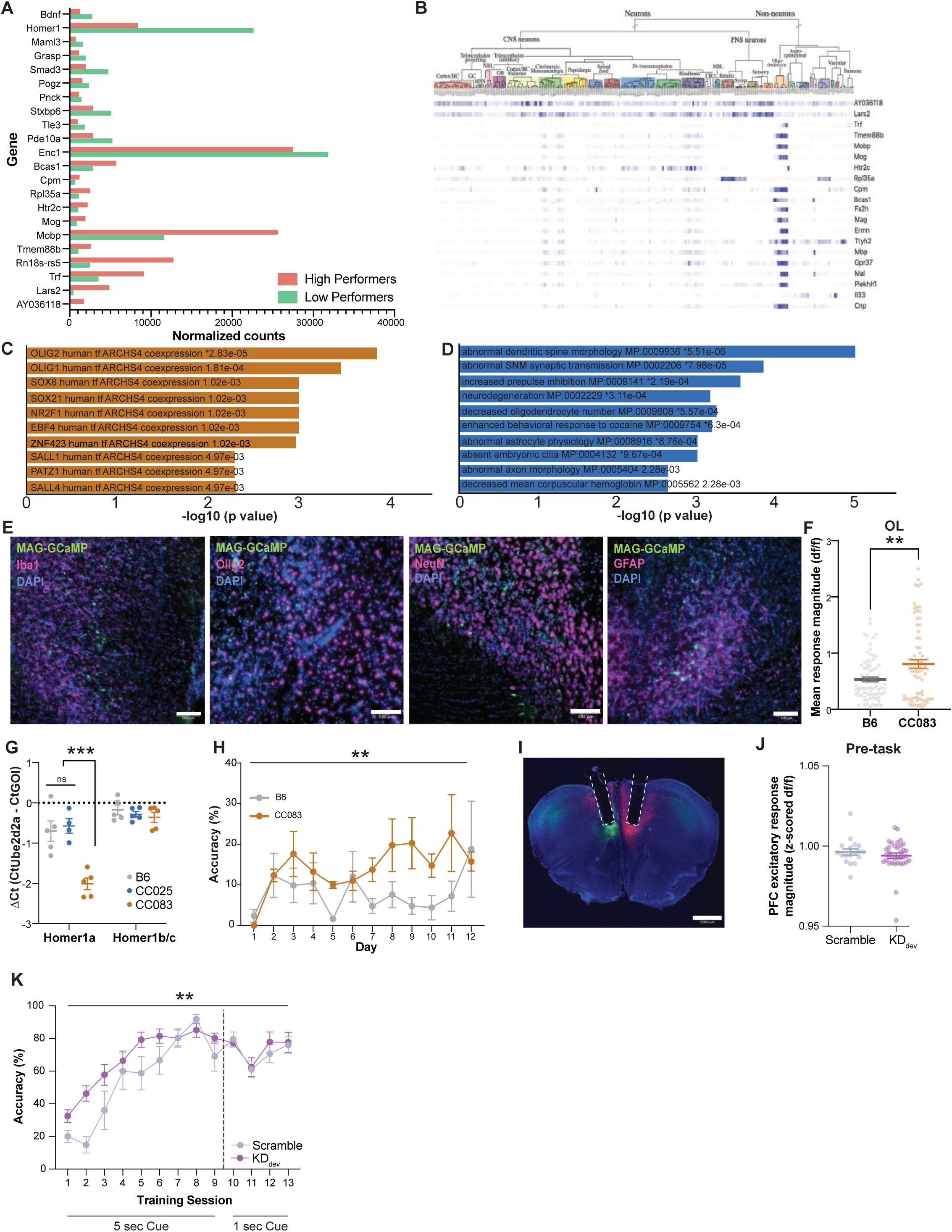
Generation and validation of MAG-GCaMP6f AAV construct, Homer1 isoform expression, and SDT behavioral performance, related to Figure 6. (**A**) The most differentially expressed genes between DO high-(n=3) and DO low-(n=3) performers from bulk PFC RNAseq (Fig. 3b). Genes are ordered by Log2FC relative to high-performers (11 upregulated and 11 downregulated). (**B**) Typical cell-type expression patterns for the 20 most upregulated genes in PFC of DO mice with high PPI performance. Differential gene expression between performance groups was determined by DESeq2, n=6 biologically independent samples. Cell-type expression patterns were determined by the Linnarsson Lab adult mouse single cell gene expression database (http://www.mousebrain.org). Genes selected for each group had log2 fold change ≥ 0.7 and adjusted p value ≤ 0.05. (**C**) Transcription factor co-expression enrichment analysis (from ARCHS4 database) of genes significantly upregulated in the PFC of CC083 mice (n=3) relative to CC025 mice (n=3) from bulk RNAseq data (Fig. 3c). (**D**) Mammalian phenotype ontology enrichment analysis (from MGI Mammalian Phenotype database with Level 4 cutoff) of genes upregulated in CC025 cells within neuron subcluster 0 from scRNAseq (Fig. 5c). (**E**) Immunohistochemistry performed 4 weeks after PFC injections of AAV-MAG-GCaMP6f (green) showing viral transduction in the target area. 20x images were collected of sections stained with antibodies (magenta) raised against the microglial marker Iba1 (top left), neuronal marker NeuN (bottom left), oligodendrocytes lineage marker Olig2 (top right), and astrocytic marker GFAP (bottom right), as well as DAPI (blue). (**F**) Average activity (area under responses) in home cage for B6 (n=5) vs CC083 (n=4) during 1 min recordings from PFC oligodendrocytes (OLs, Welch-corrected t-test p=0.004). (**G**) PFC expression of *Homer1a* and *Homer1b/c* by qPCR in B6, CC025, and CC083 adult mice (*Homer1a*: n(B6)=5, n(CC025)=4, and n(CC083)=5; *Homer1b/c:* n=5 per line; two-way ANOVA showed significant main effects for strain, *Homer1* isoform, and a significant interaction between those variables, p<0.001 for all; post-hoc Holm-Sidak’s test showed significant differences for B6 *vs* CC083 and CC025 vs CC083, p<0.001 for both). (**H**) Performance of B6 (n=5) and CC083 (n=4) mice during SDT across days showing the percentage of correct responses (two-way ANOVA, p=0.002). Tethering mice to fibers impacted performance for both lines equally. (**I**) Representative DAPI-stained (blue) histology image of dual-color photometry surgical preparation to simultaneously record from excitatory and inhibitory neurons in PFC by injecting AAV-mDlx-GCaMP6f (green) contralateral to AAV-CaMKII-Cre + AAV-CAG-FLEX-jRGECO1a (red) and implanting fibers above the injection site (indicated by white dashed outlines). (**J**) Average activity (area under responses) in home cage for Scramble (n=6) vs KD_dev_ (n=10) during 1 min recordings from PFC excitatory neurons. (**K**) Accuracy (percentage of correct responses) for Scramble (n=6) and KD_dev_ mice (n=10). Two way ANOVA showed a significant interaction between training session and group (p=0.002).

## REFERENCES

1. Bushnell, P.J. (1998). Behavioral approaches to the assessment of attention in animals. Psychopharmacology 138, 231–259. 10.1007/s002130050668.

2. Sarter, M., Givens, B., and Bruno, J.P. (2001). The cognitive neuroscience of sustained attention: where top-down meets bottom-up. Brain Research Reviews 35, 146–160. 10.1016/S0165-0173(01)00044-3.

3. Dayan, P., Kakade, S., and Montague, P.R. (2000). Learning and selective attention. Nat Neurosci 3, 1218–1223. 10.1038/81504.

4. Robbins, T.W. (1997). Arousal systems and attentional processes. Biol Psychol 45, 57–71. 10.1016/s0301-0511(96)05222-2.

5. Petersen, S.E., and Posner, M.I. (2012). The attention system of the human brain: 20 years after. Annu Rev Neurosci 35, 73–89. 10.1146/annurev-neuro-062111-150525.

6. Harris, K.D., and Thiele, A. (2011). Cortical state and attention. Nat Rev Neurosci 12, 509–523. 10.1038/nrn3084.

7. Arnsten, A.F., Wang, M.J., and Paspalas, C.D. (2012). Neuromodulation of thought: flexibilities and vulnerabilities in prefrontal cortical network synapses. Neuron 76, 223–239. 10.1016/j.neuron.2012.08.038.

8. Buschman, T.J., and Kastner, S. (2015). From Behavior to Neural Dynamics: An Integrated Theory of Attention. Neuron 88, 127–144. 10.1016/j.neuron.2015.09.017.

9. Halassa, M.M., and Kastner, S. (2017). Thalamic functions in distributed cognitive control. Nat Neurosci 20, 1669–1679. 10.1038/s41593-017-0020-1.

10. Varela, C. (2014). Thalamic neuromodulation and its implications for executive networks. Front Neural Circuits 8, 69. 10.3389/fncir.2014.00069.

11. Lee, S.H., and Dan, Y. (2012). Neuromodulation of brain states. Neuron 76, 209–222. 10.1016/j.neuron.2012.09.012.

12. Aston-Jones, G., and Cohen, J.D. (2005). An integrative theory of locus coeruleus-norepinephrine function: adaptive gain and optimal performance. Annu Rev Neurosci 28, 403–450. 10.1146/annurev.neuro.28.061604.135709.

13. Robbins, T.W., and Arnsten, A.F. (2009). The neuropsychopharmacology of fronto-executive function: monoaminergic modulation. Annu Rev Neurosci 32, 267–287. 10.1146/annurev.neuro.051508.135535.

14. Bargmann, C.I., and Marder, E. (2013). From the connectome to brain function. Nat Methods 10, 483–490. 10.1038/nmeth.2451.

15. Thiele, A., and Bellgrove, M.A. (2018). Neuromodulation of Attention. Neuron 97, 769–785. 10.1016/j.neuron.2018.01.008.

16. Chappell, P.B., Riddle, M.A., Scahill, L., Lynch, K.A., Schultz, R., Arnsten, A., Leckman, J.F., and Cohen, D.J. (1995). Guanfacine treatment of comorbid attention-deficit hyperactivity disorder and Tourette’s syndrome: preliminary clinical experience. J Am Acad Child Adolesc Psychiatry 34, 1140–1146. 10.1097/00004583-199509000-00010.

17. Dudai, Y., Jan, Y.N., Byers, D., Quinn, W.G., and Benzer, S. (1976). dunce, a mutant of Drosophila deficient in learning. Proc Natl Acad Sci U S A 73, 1684–1688. 10.1073/pnas.73.5.1684.

18. Bargiello, T.A., Jackson, F.R., and Young, M.W. (1984). Restoration of circadian behavioural rhythms by gene transfer in Drosophila. Nature 312, 752–754. 10.1038/312752a0.

19. 19. de Bono, M., and Bargmann, C.I. (1998). Natural variation in a neuropeptide Y receptor homolog modifies social behavior and food response in C. elegans. Cell 94, 679–689. 10.1016/s0092-8674(00)81609-8.

20. Bendesky, A., Kwon, Y.M., Lassance, J.M., Lewarch, C.L., Yao, S., Peterson, B.K., He, M.X., Dulac, C., and Hoekstra, H.E. (2017). The genetic basis of parental care evolution in monogamous mice. Nature 544, 434–439. 10.1038/nature22074.

21. Hsiao, K., Noble, C., Pitman, W., Yadav, N., Kumar, S., Keele, G.R., Terceros, A., Kanke, M., Conniff, T., Cheleuitte-Nieves, C., et al. (2020). A Thalamic Orphan Receptor Drives Variability in Short-Term Memory. Cell 183, 522–536.e19. 10.1016/j.cell.2020.09.011.

22. Blumenthal, T.D. (2015). Presidential Address 2014: The more-or-less interrupting effects of the startle response. Psychophysiology 52, 1417–1431. 10.1111/psyp.12506.

23. Geyer, M.A. (2006). The family of sensorimotor gating disorders: Comorbidities or diagnostic overlaps? neurotox res 10, 211–220. 10.1007/BF03033358.

24. Fendt, M., Li, L., and Yeomans, J.S. (2001). Brain stem circuits mediating prepulse inhibition of the startle reflex. Psychopharmacology 156, 216–224. 10.1007/s002130100794.

25. Vales, K., and Holubova, K. (2021). Minireview: Animal model of schizophrenia from the perspective of behavioral pharmacology: Effect of treatment on cognitive functions. Neuroscience Letters 761, 136098. 10.1016/j.neulet.2021.136098.

26. Ellenbroek, Bart A. (2004). Pre-attentive processing and schizophrenia: animal studies. Psychopharmacology 174. 10.1007/s00213-003-1684-7.

27. Green, M.F., Butler, P.D., Chen, Y., Geyer, M.A., Silverstein, S., Wynn, J.K., Yoon, J.H., and Zemon, V. (2009). Perception Measurement in Clinical Trials of Schizophrenia: Promising Paradigms From CNTRICS. Schizophrenia Bulletin 35, 163–181. 10.1093/schbul/sbn156.

28. Koch, M., and Schnitzler, H.U. (1997). The acoustic startle response in rats–circuits mediating evocation, inhibition and potentiation. Behav Brain Res 89, 35–49. 10.1016/s0166-4328(97)02296-1.

29. Braff, D.L., Geyer, M.A., and Swerdlow, N.R. (2001). Human studies of prepulse inhibition of startle: normal subjects, patient groups, and pharmacological studies. Psychopharmacology 156, 234–258. 10.1007/s002130100810.

30. Oliveras, I., Río-Álamos, C., Cañete, T., Blázquez, G., Martínez-Membrives, E., Giorgi, O., Corda, M.G., Tobeña, A., and Fernández-Teruel, A. (2015). Prepulse inhibition predicts spatial working memory performance in the inbred Roman high- and low-avoidance rats and in genetically heterogeneous NIH-HS rats: relevance for studying pre-attentive and cognitive anomalies in schizophrenia. Front. Behav. Neurosci. 9. 10.3389/fnbeh.2015.00213.

31. Ding, Y., Tian, Q., Hou, W., Chen, Z., Mao, Z., Bo, Q., Dong, F., and Wang, C. (2023). Core of sensory gating deficits in first-episode schizophrenia: attention dysfunction. Front. Psychiatry 14, 1160715. 10.3389/fpsyt.2023.1160715.

32. Li, L., Du, Y., Li, N., Wu, X., and Wu, Y. (2009). Top-down modulation of prepulse inhibition of the startle reflex in humans and rats. Neurosci Biobehav Rev 33, 1157–1167. 10.1016/j.neubiorev.2009.02.001.

33. Scholes, K.E., and Martin-Iverson, M.T. (2009). Relationships between prepulse inhibition and cognition are mediated by attentional processes. Behav Brain Res 205, 456–467. 10.1016/j.bbr.2009.07.031.

34. Scholes, K.E., and Martin-Iverson, M.T. (2010). Disturbed prepulse inhibition in patients with schizophrenia is consequential to dysfunction of selective attention. Psychophysiology 47, 223–235. 10.1111/j.1469-8986.2009.00927.x.

35. Broman, K.W., Gatti, D.M., Simecek, P., Furlotte, N.A., Prins, P., Sen, S., Yandell, B.S., and Churchill, G.A. (2019). R/qtl2: Software for Mapping Quantitative Trait Loci with High-Dimensional Data and Multiparent Populations. Genetics 211, 495–502. 10.1534/genetics.118.301595.

36. Turner, K.M., Peak, J., and Burne, T.H.J. (2016). Measuring Attention in Rodents: Comparison of a Modified Signal Detection Task and the 5-Choice Serial Reaction Time Task. Frontiers in Behavioral Neuroscience 9, 370. 10.3389/fnbeh.2015.00370.

37. Bushnell, P.J., and Strupp, B.J. (2009). Assessing Attention in Rodents. In Methods of Behavior Analysis in Neuroscience Frontiers in Neuroscience., J. J. Buccafusco, ed. (CRC Press/Taylor & Francis).

38. Callahan, P.M., and Terry, A.V. (2015). Attention. Handb Exp Pharmacol 228, 161–189. 10.1007/978-3-319-16522-6_5.

39. Bottai, D., Guzowski, J.F., Schwarz, M.K., Kang, S.H., Xiao, B., Lanahan, A., Worley, P.F., and Seeburg, P.H. (2002). Synaptic Activity-Induced Conversion of Intronic to Exonic Sequence in Homer 1 Immediate Early Gene Expression. The Journal of Neuroscience 22, 167. 10.1523/JNEUROSCI.22-01-00167.2002.

40. Brakeman, P.R., Lanahan, A.A., O’Brien, R., Roche, K., Barnes, C.A., Huganir, R.L., and Worley, P.F. (1997). Homer: a protein that selectively binds metabotropic glutamate receptors. Nature 386, 284–288. 10.1038/386284a0.

41. Kato, A., Ozawa, F., Saitoh, Y., Hirai, K., and Inokuchi, K. (1997). vesl, a gene encoding VASP/Ena family related protein, is upregulated during seizure, long-term potentiation and synaptogenesis. FEBS Lett 412, 183–189. 10.1016/s0014-5793(97)00775-8.

42. Xiao, B., Tu, J.C., Petralia, R.S., Yuan, J.P., Doan, A., Breder, C.D., Ruggiero, A., Lanahan, A.A., Wenthold, R.J., and Worley, P.F. (1998). Homer Regulates the Association of Group 1 Metabotropic Glutamate Receptors with Multivalent Complexes of Homer-Related, Synaptic Proteins. Neuron 21, 707–716. 10.1016/S0896-6273(00)80588-7.

43. Szumlinski, K.K., Lominac, K.D., Kleschen, M.J., Oleson, E.B., Dehoff, M.H., Schwarz, M.K., Seeburg, P.H., Worley, P.F., and Kalivas, P.W. (2005). Behavioral and neurochemical phenotyping of Homer1 mutant mice: possible relevance to schizophrenia. Genes Brain Behav 4, 273–288. 10.1111/j.1601-183X.2005.00120.x.

44. Lominac, K.D., Oleson, E.B., Pava, M., Klugmann, M., Schwarz, M.K., Seeburg, P.H., During, M.J., Worley, P.F., Kalivas, P.W., and Szumlinski, K.K. (2005). Distinct Roles for Different Homer1 Isoforms in Behaviors and Associated Prefrontal Cortex Function. The Journal of Neuroscience 25, 11586. 10.1523/JNEUROSCI.3764-05.2005.

45. Jaubert, P.J., Golub, M.S., Lo, Y.Y., Germann, S.L., Dehoff, M.H., Worley, P.F., Kang, S.H., Schwarz, M.K., Seeburg, P.H., and Berman, R.F. (2006). Complex, multimodal behavioral profile of the Homer1 knockout mouse. Genes, Brain and Behavior 6, 141–154. 10.1111/j.1601-183X.2006.00240.x.

46. Datko, M.C., Hu, J.-H., Williams, M., Reyes, C.M., Lominac, K.D., von Jonquieres, G., Klugmann, M., Worley, P.F., and Szumlinski, K.K. (2017). Behavioral and Neurochemical Phenotyping of Mice Incapable of Homer1a Induction. Frontiers in Behavioral Neuroscience 11, 208. 10.3389/fnbeh.2017.00208.

47. Muzzio, I.A., Levita, L., Kulkarni, J., Monaco, J., Kentros, C., Stead, M., Abbott, L.F., and Kandel, E.R. (2009). Attention Enhances the Retrieval and Stability of Visuospatial and Olfactory Representations in the Dorsal Hippocampus. PLoS Biol 7, e1000140. 10.1371/journal.pbio.1000140.

48. Zeisel, A., Hochgerner, H., Lönnerberg, P., Johnsson, A., Memic, F., van der Zwan, J., Häring, M., Braun, E., Borm, L.E., La Manno, G., et al. (2018). Molecular Architecture of the Mouse Nervous System. Cell 174, 999–1014.e22. 10.1016/j.cell.2018.06.021.

49. Shiraishi-Yamaguchi, Y., and Furuichi, T. (2007). The Homer family proteins. Genome Biology 8, 206. 10.1186/gb-2007-8-2-206.

50. Ango, F., Pin, J.P., Tu, J.C., Xiao, B., Worley, P.F., Bockaert, J., and Fagni, L. (2000). Dendritic and axonal targeting of type 5 metabotropic glutamate receptor is regulated by homer1 proteins and neuronal excitation. J Neurosci 20, 8710–8716. 10.1523/JNEUROSCI.20-23-08710.2000.

51. Petralia, R.S., Wang, Y.X., Sans, N., Worley, P.F., Hammer, 3rd, J.A., and Wenthold, R.J. (2001). Glutamate receptor targeting in the postsynaptic spine involves mechanisms that are independent of myosin Va. Eur J Neurosci 13, 1722–1732. 10.1046/j.0953-816x.2001.01553.x.

52. Finak, G., McDavid, A., Yajima, M., Deng, J., Gersuk, V., Shalek, A.K., Slichter, C.K., Miller, H.W., McElrath, M.J., Prlic, M., et al. (2015). MAST: a flexible statistical framework for assessing transcriptional changes and characterizing heterogeneity in single-cell RNA sequencing data. Genome Biology 16, 278. 10.1186/s13059-015-0844-5.

53. Chen, E.Y., Tan, C.M., Kou, Y., Duan, Q., Wang, Z., Meirelles, G.V., Clark, N.R., and Ma’ayan, A. (2013). Enrichr: interactive and collaborative HTML5 gene list enrichment analysis tool. BMC Bioinformatics 14, 128. 10.1186/1471-2105-14-128.

54. Kuleshov, M.V., Jones, M.R., Rouillard, A.D., Fernandez, N.F., Duan, Q., Wang, Z., Koplev, S., Jenkins, S.L., Jagodnik, K.M., Lachmann, A., et al. (2016). Enrichr: a comprehensive gene set enrichment analysis web server 2016 update. Nucleic Acids Res 44, W90–7. 10.1093/nar/gkw377.

55. Xie, Z., Bailey, A., Kuleshov, M.V., Clarke, D.J.B., Evangelista, J.E., Jenkins, S.L., Lachmann, A., Wojciechowicz, M.L., Kropiwnicki, E., Jagodnik, K.M., et al. (2021). Gene Set Knowledge Discovery with Enrichr. Current Protocols 1, e90. 10.1002/cpz1.90.

56. de Lecea, L., Carter, M.E., and Adamantidis, A. (2012). Shining light on wakefulness and arousal. Biol Psychiatry 71, 1046–1052. 10.1016/j.biopsych.2012.01.032.

57. Lovett-Barron, M., Andalman, A.S., Allen, W.E., Vesuna, S., Kauvar, I., Burns, V.M., and Deisseroth, K. (2017). Ancestral Circuits for the Coordinated Modulation of Brain State. Cell 171, 1411–1423 e17. 10.1016/j.cell.2017.10.021.

58. Banaschewski, T., Roessner, V., Dittmann, R.W., Santosh, P.J., and Rothenberger, A. (2004). Non-stimulant medications in the treatment of ADHD. Eur Child Adolesc Psychiatry 13 Suppl 1, I102–16. 10.1007/s00787-004-1010-x.

59. Cinnamon Bidwell, L., Dew, R.E., and Kollins, S.H. (2010). Alpha-2 adrenergic receptors and attention-deficit/hyperactivity disorder. Curr Psychiatry Rep 12, 366–373. 10.1007/s11920-010-0136-4.

60. Elia, J., Glessner, J.T., Wang, K., Takahashi, N., Shtir, C.J., Hadley, D., Sleiman, P.M.A., Zhang, H., Kim, C.E., Robison, R., et al. (2012). Genome-wide copy number variation study associates metabotropic glutamate receptor gene networks with attention deficit hyperactivity disorder. Nat Genet 44, 78–84. 10.1038/ng.1013.

61. Sánchez-Mora, C., Soler Artigas, M., Garcia-Martínez, I., Pagerols, M., Rovira, P., Richarte, V., Corrales, M., Fadeuilhe, C., Padilla, N., De La Cruz, X., et al. (2019). Epigenetic signature for attention-deficit/hyperactivity disorder: identification of miR-26b-5p, miR-185-5p, and miR-191-5p as potential biomarkers in peripheral blood mononuclear cells. Neuropsychopharmacol. 44, 890–897. 10.1038/s41386-018-0297-0.

62. Hong, Q., Zhang, M., Pan, X.Q., Guo, M., Li, F., Tong, M.L., Chen, R.H., Guo, X.R., and Chi, X. (2009). Prefrontal cortex Homer expression in an animal model of attention-deficit/hyperactivity disorder. J Neurol Sci 287, 205–211. 10.1016/j.jns.2009.07.024.

63. Naaijen, J., Bralten, J., Poelmans, G., Consortium, I., Glennon, J.C., Franke, B., and Buitelaar, J.K. (2017). Glutamatergic and GABAergic gene sets in attention-deficit/hyperactivity disorder: association to overlapping traits in ADHD and autism. Transl Psychiatry 7, e999. 10.1038/tp.2016.273.

64. Norton, N., Williams, H.J., Williams, N.M., Spurlock, G., Zammit, S., Jones, G., Jones, S., Owen, R., O’Donovan, M.C., and Owen, M.J. (2003). Mutation screening of the Homer gene family and association analysis in schizophrenia. Am J Med Genet B Neuropsychiatr Genet 120B, 18–21. 10.1002/ajmg.b.20032.

65. Spellmann, I., Rujescu, D., Musil, R., Mayr, A., Giegling, I., Genius, J., Zill, P., Dehning, S., Opgen-Rhein, M., Cerovecki, A., et al. (2011). Homer-1 polymorphisms are associated with psychopathology and response to treatment in schizophrenic patients. J Psychiatr Res 45, 234–241. 10.1016/j.jpsychires.2010.06.004.

66. Kelleher, 3rd, R.J., Geigenmuller, U., Hovhannisyan, H., Trautman, E., Pinard, R., Rathmell, B., Carpenter, R., and Margulies, D. (2012). High-throughput sequencing of mGluR signaling pathway genes reveals enrichment of rare variants in autism. PLoS One 7, e35003. 10.1371/journal.pone.0035003.

67. Gai, X., Xie, H.M., Perin, J.C., Takahashi, N., Murphy, K., Wenocur, A.S., D’Arcy, M., O’Hara, R.J., Goldmuntz, E., Grice, D.E., et al. (2012). Rare structural variation of synapse and neurotransmission genes in autism. Mol Psychiatry 17, 402–411. 10.1038/mp.2011.10.

68. Sala, C., Futai, K., Yamamoto, K., Worley, P.F., Hayashi, Y., and Sheng, M. (2003). Inhibition of dendritic spine morphogenesis and synaptic transmission by activity-inducible protein Homer1a. J Neurosci 23, 6327–6337. 10.1523/JNEUROSCI.23-15-06327.2003.

69. Flavell, S.W., Kim, T.K., Gray, J.M., Harmin, D.A., Hemberg, M., Hong, E.J., Markenscoff-Papadimitriou, E., Bear, D.M., and Greenberg, M.E. (2008). Genome-wide analysis of MEF2 transcriptional program reveals synaptic target genes and neuronal activity-dependent polyadenylation site selection. Neuron 60, 1022–1038. 10.1016/j.neuron.2008.11.029.

70. Diering, G.H., Nirujogi, R.S., Roth, R.H., Worley, P.F., Pandey, A., and Huganir, R.L. (2017). Homer1a drives homeostatic scaling-down of excitatory synapses during sleep. Science 355, 511–515. 10.1126/science.aai8355.

71. Bockaert, J., Perroy, J., and Ango, F. (2021). The Complex Formed by Group I Metabotropic Glutamate Receptor (mGluR) and Homer1a Plays a Central Role in Metaplasticity and Homeostatic Synaptic Scaling. J Neurosci 41, 5567–5578. 10.1523/JNEUROSCI.0026-21.2021.

72. Gibson, E.M., Purger, D., Mount, C.W., Goldstein, A.K., Lin, G.L., Wood, L.S., Inema, I., Miller, S.E., Bieri, G., Zuchero, J.B., et al. (2014). Neuronal activity promotes oligodendrogenesis and adaptive myelination in the mammalian brain. Science 344, 1252304. 10.1126/science.1252304.

73. Hughes, E.G., Orthmann-Murphy, J.L., Langseth, A.J., and Bergles, D.E. (2018). Myelin remodeling through experience-dependent oligodendrogenesis in the adult somatosensory cortex. Nature Neuroscience 21, 696–706. 10.1038/s41593-018-0121-5.

74. Geraghty, A.C., Gibson, E.M., Ghanem, R.A., Greene, J.J., Ocampo, A., Goldstein, A.K., Ni, L., Yang, T., Marton, R.M., Paşca, S.P., et al. (2019). Loss of Adaptive Myelination Contributes to Methotrexate Chemotherapy-Related Cognitive Impairment. Neuron 103, 250–265.e8. 10.1016/j.neuron.2019.04.032.

75. Noori, R., Park, D., Griffiths, J.D., Bells, S., Frankland, P.W., Mabbott, D., and Lefebvre, J. (2020). Activity-dependent myelination: A glial mechanism of oscillatory self-organization in large-scale brain networks. Proceedings of the National Academy of Sciences 117, 13227– 13237. 10.1073/pnas.1916646117.

76. Che, A., Babij, R., Iannone, A.F., Fetcho, R.N., Ferrer, M., Liston, C., Fishell, G., and De Marco Garcia, N.V. (2018). Layer I Interneurons Sharpen Sensory Maps during Neonatal Development. Neuron 99, 98–116 e7. 10.1016/j.neuron.2018.06.002.

77. Dana, H., Mohar, B., Sun, Y., Narayan, S., Gordus, A., Hasseman, J.P., Tsegaye, G., Holt, G.T., Hu, A., Walpita, D., et al. (2016). Sensitive red protein calcium indicators for imaging neural activity. Elife 5. 10.7554/eLife.12727.

78. Chen, T.W., Wardill, T.J., Sun, Y., Pulver, S.R., Renninger, S.L., Baohan, A., Schreiter, E.R., Kerr, R.A., Orger, M.B., Jayaraman, V., et al. (2013). Ultrasensitive fluorescent proteins for imaging neuronal activity. Nature 499, 295–300. 10.1038/nature12354.

79. Powell, S.K., Khan, N., Parker, C.L., Samulski, R.J., Matsushima, G., Gray, S.J., and McCown, T.J. (2016). Characterization of a novel adeno-associated viral vector with preferential oligodendrocyte tropism. Gene Ther 23, 807–814. 10.1038/gt.2016.62.

80. Jin, J., Cheng, J., Lee, K.-W., Amreen, B., McCabe, K.A., Pitcher, C., Liebmann, T., Greengard, P., and Flajolet, M. (2019). Cholinergic Neurons of the Medial Septum Are Crucial for Sensorimotor Gating. The Journal of Neuroscience 39, 5234. 10.1523/JNEUROSCI.0950-18.2019.

81. Lister, R.G. (1987). The use of a plus-maze to measure anxiety in the mouse. Psychopharmacology (Berl) 92, 180–185. 10.1007/BF00177912.

82. Kang, H.M., Sul, J.H., Service, S.K., Zaitlen, N.A., Kong, S.Y., Freimer, N.B., Sabatti, C., and Eskin, E. (2010). Variance component model to account for sample structure in genome-wide association studies. Nat Genet 42, 348–354. 10.1038/ng.548.

83. Azizi, E., Carr, A.J., Plitas, G., Cornish, A.E., Konopacki, C., Prabhakaran, S., Nainys, J., Wu, K., Kiseliovas, V., Setty, M., et al. (2018). Single-Cell Map of Diverse Immune Phenotypes in the Breast Tumor Microenvironment. Cell 174, 1293–1308 e36. 10.1016/j.cell.2018.05.060.

84. Wolock, S.L., Lopez, R., and Klein, A.M. (2019). Scrublet: Computational Identification of Cell Doublets in Single-Cell Transcriptomic Data. Cell Syst 8, 281–291 e9. 10.1016/j.cels.2018.11.005.

85. von Jonquieres, G., Frohlich, D., Klugmann, C.B., Wen, X., Harasta, A.E., Ramkumar, R., Spencer, Z.H., Housley, G.D., and Klugmann, M. (2016). Recombinant Human Myelin-Associated Glycoprotein Promoter Drives Selective AAV-Mediated Transgene Expression in Oligodendrocytes. Front Mol Neurosci 9, 13. 10.3389/fnmol.2016.00013.

86. Frazer, K.A., Pachter, L., Poliakov, A., Rubin, E.M., and Dubchak, I. (2004). VISTA: computational tools for comparative genomics. Nucleic Acids Res 32, W273–9. 10.1093/nar/gkh458.

87. Hu, S., Xie, Z., Onishi, A., Yu, X., Jiang, L., Lin, J., Rho, H.S., Woodard, C., Wang, H., Jeong, J.S., et al. (2009). Profiling the human protein-DNA interactome reveals ERK2 as a transcriptional repressor of interferon signaling. Cell 139, 610–622. 10.1016/j.cell.2009.08.037.

88. Zhang, Z., Wang, W., and Valdar, W. (2014). Bayesian modeling of haplotype effects in multiparent populations. Genetics 198, 139–156. 10.1534/genetics.114.166249.

89. Love, M.I., Huber, W., and Anders, S. (2014). Moderated estimation of fold change and dispersion for RNA-seq data with DESeq2. Genome Biology 15, 550. 10.1186/s13059-014-0550-8.

90. Dobin, A., Davis, C.A., Schlesinger, F., Drenkow, J., Zaleski, C., Jha, S., Batut, P., Chaisson, M., and Gingeras, T.R. (2013). STAR: ultrafast universal RNA-seq aligner. Bioinformatics 29, 15–21. 10.1093/bioinformatics/bts635.

91. Patro, R., Duggal, G., Love, M.I., Irizarry, R.A., and Kingsford, C. (2017). Salmon provides fast and bias-aware quantification of transcript expression. Nature Methods 14, 417–419. 10.1038/nmeth.4197.

92. Wolf, F.A., Angerer, P., and Theis, F.J. (2018). SCANPY: large-scale single-cell gene expression data analysis. Genome Biology 19, 15. 10.1186/s13059-017-1382-0.

